# Lactate transport inhibition therapeutically reprograms fibroblast metabolism in experimental pulmonary fibrosis

**DOI:** 10.1101/2024.04.25.591150

**Authors:** David R. Ziehr, Fei Li, K. Mark Parnell, Nathan M. Krah, Kevin J. Leahy, Christelle Guillermier, Jack Varon, Rebecca M. Baron, Bradley A. Maron, Nancy J. Philp, Lida P. Hariri, Edy Y. Kim, Matthew L. Steinhauser, Rachel S. Knipe, Jared Rutter, William M. Oldham

## Abstract

Myofibroblast differentiation, essential for driving extracellular matrix synthesis in pulmonary fibrosis, requires increased glycolysis. While glycolytic cells must export lactate, the contributions of lactate transporters to myofibroblast differentiation are unknown. In this study, we investigated how MCT1 and MCT4, key lactate transporters, influence myofibroblast differentiation and experimental pulmonary fibrosis. Our findings reveal that inhibiting MCT1 or MCT4 reduces TGFβ-stimulated pulmonary myofibroblast differentiation *in vitro* and decreases bleomycin-induced pulmonary fibrosis *in vivo*. Through comprehensive metabolic analyses, including bioenergetics, stable isotope tracing, metabolomics, and imaging mass spectrometry in both cells and mice, we demonstrate that inhibiting lactate transport enhances oxidative phosphorylation, reduces reactive oxygen species production, and diminishes glucose metabolite incorporation into fibrotic lung regions. Furthermore, we introduce VB253, a novel MCT4 inhibitor, which ameliorates pulmonary fibrosis in both young and aged mice, with comparable efficacy to established antifibrotic therapies. These results underscore the necessity of lactate transport for myofibroblast differentiation, identify MCT1 and MCT4 as promising pharmacologic targets in pulmonary fibrosis, and support further evaluation of lactate transport inhibitors for patients for whom limited therapeutic options currently exist.

**SUMMARY:** Small molecule inhibitors of lactate transporters, including the novel MCT4 inhibitor VB253, reprogram fibroblast metabolism to prevent myofibroblast differentiation and decrease bleomycin-induced pulmonary fibrosis.

## INTRODUCTION

Idiopathic pulmonary fibrosis (IPF) is a chronic and progressive lung disease with high mortality and limited therapeutic options. IPF affects approximately 150,000 patients in the U.S. with a median survival of 3-5 years (*1–3*). Currently approved pharmacotherapies for IPF are limited to the antifibrotics pirfenidone and nintedanib that slow, but do not stop, disease progression (*4, 5*), leaving lung transplantation as the only option available to eligible patients with progressive disease. The limited efficacy of antifibrotic therapies emphasizes the need for novel therapeutic approaches targeting different features of IPF pathobiology.

Accumulating evidence suggests that metabolic reprogramming may be one such therapeutic strategy in IPF (*6, 7*). Lung fibrosis is driven by the excessive deposition of extracellular matrix by myofibroblasts (*3*). Fundamental changes in myofibroblast metabolism support myofibroblast differentiation and extracellular matrix production (*8–12*). In particular, increased glycolysis and lactate production have been observed in IPF myofibroblasts *ex vivo* and following transforming growth factor β1 (TGFβ)-induced myofibroblast differentiation *in vitro* (*8, 9, 13, 14*). These metabolic changes are critical for fibrogenesis, as small molecule inhibitors of glucose uptake, glycolysis, and lactate fermentation prevent myofibroblast differentiation *in vitro* and attenuate pulmonary fibrosis in animal models (*8–10, 13, 15, 16*). Unfortunately, low target affinities, poor specificity, narrow therapeutic indices, and common genetic resistance have all hampered the translation of these investigational compounds for clinical use (*17–19*). Moreover, the molecular mechanisms by which these metabolic inhibitors attenuate the myofibroblast differentiation transcriptional program remain unclear. In order to leverage metabolic therapies for IPF, more targeted and better characterized drugs must be developed.

Toward this end, we aimed to examine the impact of a novel metabolic strategy — lactate transport inhibition — on myofibroblast differentiation and experimental pulmonary fibrosis. Sustained glycolysis in myofibroblasts relies on lactate secretion, which is conducted by a family of monocarboxylate transporters (MCT1-4). Inhibitors targeting these transporters have been actively explored in clinical trials for oncological conditions where glycolytic reprogramming also features prominently in disease pathobiology (*20, 21*). Importantly, MCT inhibitors present favorable pharmacologic profiles compared to previously studied glycolysis inhibitors, with successful translation to human clinical trials for advanced solid tumors (*22*). Before the promise of this therapeutic approach in IPF may be realized, however, the preclinical efficacy and molecular mechanisms-of-action of lactate transport inhibitors must be demonstrated experimentally.

In this work, we evaluated the contribution of lactate transporters to experimental pulmonary fibrosis. We found increased expression of the lactate transporters MCT1 and MCT4 IPF patient lungs. Inhibition of these transporters attenuated bleomycin-induced lung fibrosis *in vivo* and TGFβ-induced myofibroblast differentiation *in vitro*, where MCT4 inhibition demonstrated increased therapeutic efficacy. Using metabolomics, stable isotope tracing, and high-resolution spatial metabolomic imaging, we find that lactate transport inhibition promotes glucose oxidation and decreases pro-fibrogenic reactive oxygen species (ROS) production. Building on these results, we introduce VB253, a novel MCT4 inhibitor suitable for human clinical studies that performs similarly to current standard-of-care antifibrotic therapies in bleomycin-induced pulmonary fibrosis. Together, our findings offer insights into disrupting the metabolic pathways driving IPF fibrosis and present a promising strategy targeting lactate transporters for the treatment of this fatal condition.

## RESULTS

### MCT expression increases in human pulmonary fibrosis and experimental models

Among the four lactate transporters, MCT1 and MCT4 exhibit the highest expression levels in the lung (*23*). Based on this observation, we investigated the expression of MCT1 and MCT4 in lung explants from patients with IPF obtained during transplantation. Consistent with a pathologic role for MCT1 and MCT4 in IPF, we observed a significant upregulation of both MCT1 and MCT4 proteins in IPF lung tissues compared to non-fibrotic controls (**Fig. 1A**). These findings were corroborated in an experimental model of pulmonary fibrosis, where intratracheal bleomycin administration led to increased expression of both MCT1 and MCT4 (**Fig. 1B**).

**Fig. 1.**
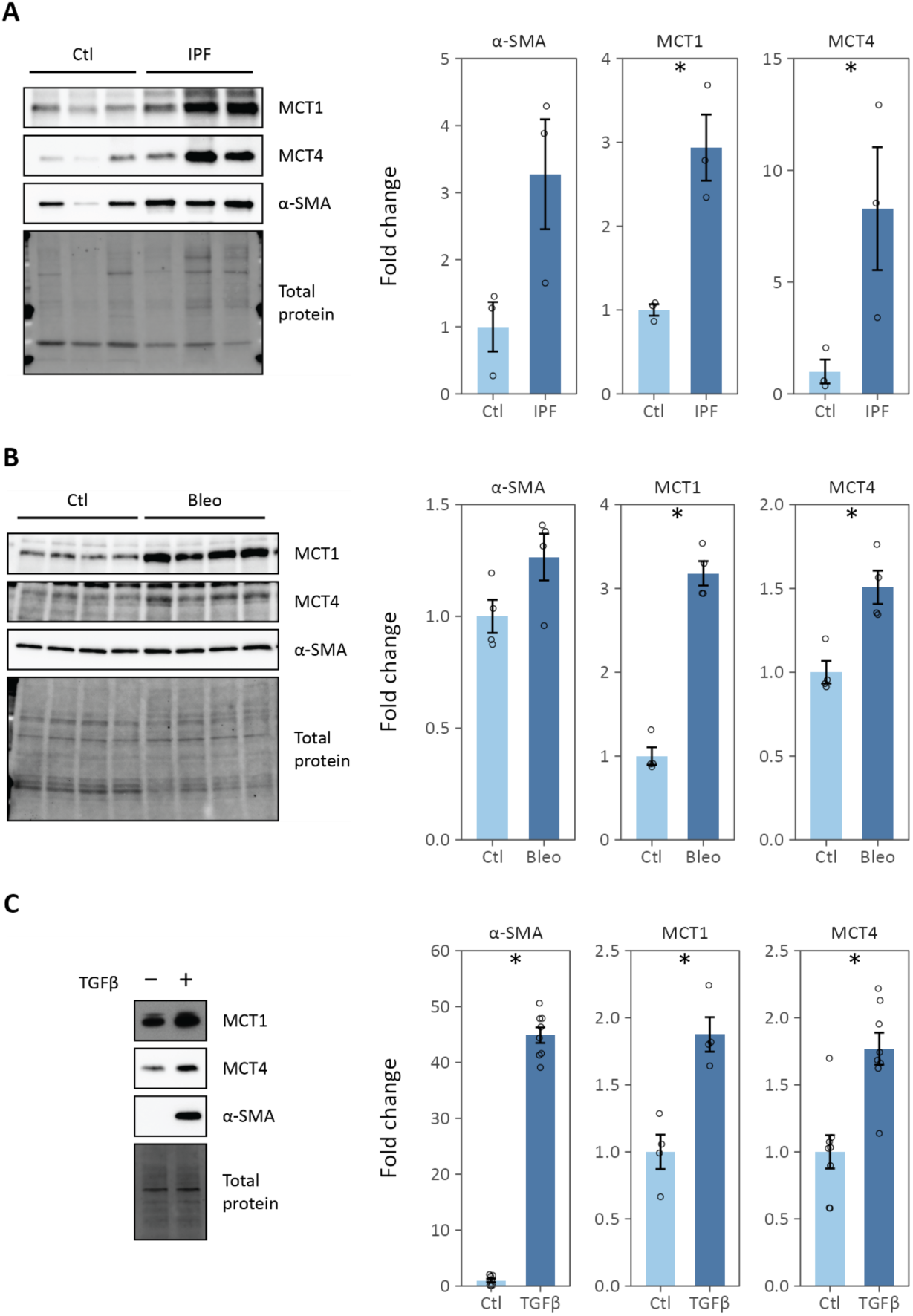
Lactate transporter expression increases in human IPF lung and experimental models. (**A**) MCT1, MCT4, and α-SMA protein expression in whole lung homogenates from explanted IPF lungs and controls (Ctl). (**B**) MCT1, MCT4, and α-SMA protein expression in whole lung homogenates from bleomycin (Bleo)- and vehicle (Ctl)-treated mice. (**C**) MCT1, MCT4, and α-SMA protein expression in cell lysates from normal human lung fibroblasts treated with TGFβ to induce myofibroblast differentiation. Individual data points are biological replicates. Summary data are mean ± SEM (* p-value < 0.05).

A hallmark of IPF is the activation of tissue myofibroblasts, distinguished by their *de novo* expression of smooth muscle α-actin (α-SMA); stress fiber formation; and increased migration, contraction, and extracellular matrix production (*7, 9, 24*). TGFβ is the most potent inducer of myofibroblast activation *in vitro* and *in vivo*. The TGFβ- dependent upregulation of α-SMA expression serves as a well-established and widely utilized model for studying myofibroblast activation pertinent to pulmonary fibrosis (*10, 25–27*). Consistent with our findings in human IPF lungs, we observed increased expression of MCT1 and MCT4 in normal human lung fibroblasts following TGFβ treatment (**Fig. 1C**). These findings align with increased expression of other glycolytic enzymes and the associated metabolic changes previously documented in these cells (*9*). Taken together, these data underscore the association between pulmonary fibrosis and lactate transporter expression in the lung generally and in myofibroblasts specifically.

### Myofibroblast differentiation *in vitro* requires lactate transport

We proceeded to investigate whether MCT expression and activity were essential for myofibroblast differentiation *in vitro* using RNA interference and pharmacologic approaches. Lung fibroblasts were transfected with siRNA targeting MCT1 and MCT4 individually and in combination. After 24 h, the cells were treated with TGFβ for 48 h to induce myofibroblast differentiation. The siRNAs reduced lactate transporter protein levels (**Fig. 2A**). Reduction in either MCT1 or MCT4 expression caused a marked decrease in TGFβ-stimulated α-SMA expression. Notably, siMCT1 also decreased MCT4 expression. No adverse effects on cell viability were noted following lactate transporter knockdown, and siMCT1 significantly increased cell count in both control and TGFβ- treated cells (**Fig. S1A**).

**Fig. 2.**
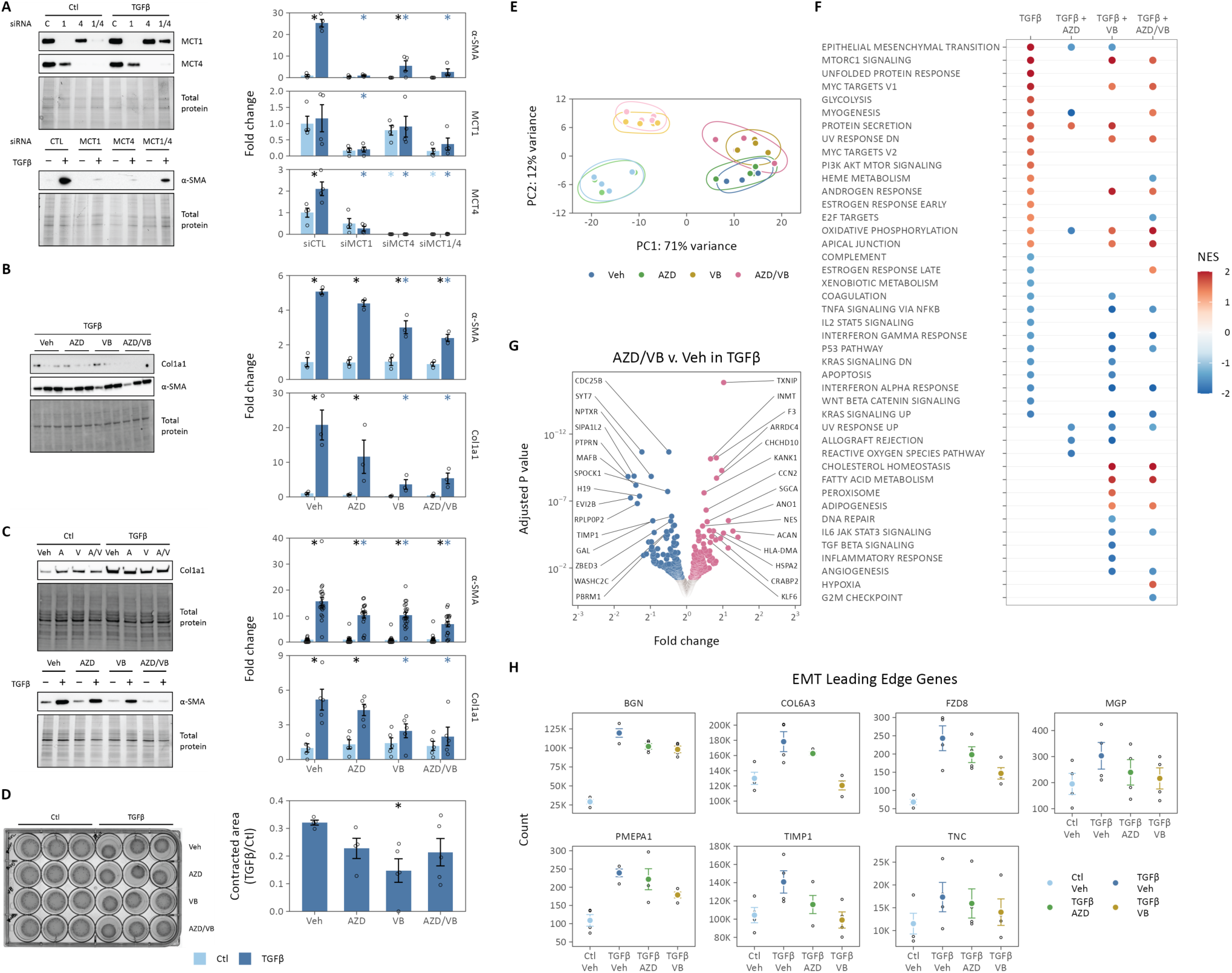
Lactate transport inhibition decreases myofibroblast differentiation and pro-fibrotic gene transcription *in vitro*. (**A**) RNA interference targeting MCT1 or MCT4 decreases TGFβ-stimulated α-SMA expression in normal human lung fibroblasts. (**B**-**C**) Small molecule inhibitors of MCT1 (AZD3965, AZD) or MCT4 (VB124, VB) decrease TGFβ-stimulated α-SMA expression in IPF lung fibroblasts (B) and normal human lung fibroblasts (C). (**D**) Lactate transport inhibitors decrease gel contractility measured 24 h following TGFβ. (**E**) Principal components analysis of RNA-seq data from lung fibroblasts treated with TGFβ (*bold colors*) or vehicle (*light colors*) and lactate transport inhibitors (N = 4). (**F**) Dot plot of Hallmark gene set enrichment analysis. Significantly enriched pathways with adjusted p-value < 0.05 are included and points are colored by normalized enrichment score (NES). Positive NES indicated relative enrichment following TGFβ compared to control (TGFβ) or with MCT inhibitor compared to vehicle in TGFβ-treated cells. (**G**) Volcano plot of significantly differentially expressed genes in TGFβ-stimulated cells treated with AZD3965 and VB124 compared to vehicle control. Significantly differentially expressed genes are highlighted (adjusted p-value < 0.05), the top 15 up- and down-regulated of which are labeled. (**H**) Transcript counts from the leading edge of enrichment for the epithelial mesenchymal transition Hallmark gene set demonstrating down- regulation of pro-fibrotic genes with MCT inhibition. Individual data points are biological replicates. Summary data are mean ± SEM (* adjusted p-value < 0.05; *black* compares TGFβ to control within a given treatment, *colored* compares the treatment effect to control for a given condition).

After observing a reduction in MCT4 expression following siMCT1 treatment, we proceeded to evaluate the impact of pharmacological MCT inhibitors on myofibroblast differentiation to examine the independent effects of MCT1 and MCT4 inhibition. AZD3965, a high-affinity (*K_i_* 1.6 nM) inhibitor of MCT1 (*28*), and VB124, a recently developed high-affinity (*K_i_* 11 nM) inhibitor of MCT4 (*21*), were used in this evaluation. IPF lung fibroblasts were differentiated with TGFβ in the presence of these MCT inhibitors (**Fig. 2B**). MCT4 inhibition by VB124 alone, or in combination with AZD3965, decreased Col1a1 and α-SMA expression. These effects were consistent with normal human lung fibroblasts where both AZD3965 and VB124, either individually or in combination, decreased α-SMA expression (**Fig. 2C**). Similarly, AR-C155858, an inhibitor with high-affinity (*K_i_* 2 nM) for both MCT1 and MCT2 (*29*), also reduced α-SMA expression, alone and in combination with VB124 (**Fig. S2B**). Importantly, pharmacologic MCT inhibition did not significantly impact cell count during the 48 h treatment (**Fig. S2C-D**). As expected, decreased Col1a1 and α-SMA expression correlated with reduced myofibroblast contractility, as demonstrated by gel contraction assay (**Fig. 2D**). Together, these data indicate that MCT expression and activity are required for myofibroblast differentiation *in vitro*.

### MCT inhibition attenuates pro-fibrotic transcriptional programs

To further characterize the antifibrotic effects of lactate transport inhibition, we conducted RNA sequencing on lung fibroblasts treated with TGFβ in conjunction with AZD3965 or VB124 (**Fig. 2E-H, S2**). Principal components analysis (PCA) revealed that the first principal component predominantly represented the effect of TGFβ treatment, while MCT4 inhibition, either alone or combined with MCT1 inhibition, aligned with the second principal component (**Fig. 2E**). Samples treated with the MCT1 inhibitor AZD3965 were similar to vehicle-treated controls.

Differential expression analysis of TGFβ-treated cells revealed the anticipated upregulation of extracellular matrix proteins (**Fig. S2A**) and enrichment of the epithelial-to-mesenchymal (EMT) gene set, among others (**Fig. 2F**). In line with the PCA results, only GRIK4 (glutamate ionotropic receptor kainate type subunit 4) and BRI3 (brain protein I3) were differentially expressed following AZD3965 treatment (**Fig. S2B**). By contrast, VB124, either alone (**Fig. S2C**) or in combination with AZD3965 (**Fig. 2G**), induced more significant alterations in fibroblast transcription, with 2% of 24,902 genes being differentially expressed at a false discovery rate (FDR) < 0.05.

Both AZD3965 and VB124 reversed TGFβ-dependent enrichment of the EMT gene set, which is the Hallmark gene set containing genes related to fibrosis (**Fig. 2F**). Leading edge analysis of the EMT gene set identified seven genes shared among all three comparisons (*i.e.*, genes increased by TGFβ and decreased by both AZD3965 and VB124) (**Fig. 2H**). These genes, including biglycan (BGN), COL6A3, Frizzled 8 (FZD8), matrix Gla protein (MGP), Prostate Transmembrane Protein, Androgen Induced 1 (PMEPA1), TIMP metallopeptidase inhibitor 1 (TIMP1), and tenascin-C (TNC), are known contributors to pulmonary fibrosis pathobiology or serve as biomarkers of disease or treatment response (*30–34*). Taken together, these findings suggest that lactate transport inhibitors attenuate the pro-fibrotic transcriptional program in TGFβ-treated lung fibroblasts.

### MCT inhibition reprograms myofibroblast metabolism

MCTs play pivotal roles in maintaining cellular lactate and redox homeostasis. MCT1 predominantly imports lactate in cells utilizing lactate for oxidative phosphorylation or gluconeogenesis and is ubiquitously expressed. MCT1 also facilitates lactate export in some glycolytic cells (*35, 36*). By contrast, MCT4 functions as the main lactate exporter in glycolytic cells and is up-regulated when the glycolytic transcriptional program is activated by, for example, the c-Myc or hypoxia-inducible transcription factors (*35, 37*). Importantly, MCT4 can also act as a lactate importer with a *K*_M_ of 1 mM (*38*).

To assess the metabolic consequences of lactate transporter inhibition, we quantified extracellular lactate in the conditioned medium from cells treated with MCT siRNA or pharmacologic inhibitors. Silencing MCT1 and MCT4, either individually or concurrently, decreased net TGFβ-stimulated lactate efflux in lung fibroblasts (**Fig. 3A**). By contrast, pharmacologic inhibition of either MCT1 or MCT4 alone did not decrease TGFβ-stimulated lactate efflux (**Fig. 3B**); inhibition of both transporters was required to lower extracellular lactate levels. Inhibition of MCT1, MCT2, and MCT4 with the combination of AR-C155858 and VB124 was required to prevent increases in lactate efflux (**Fig. S3A**). Although less pronounced compared to lactate production, we observed a trend toward decreased extracellular glucose consumption in cells treated with both AZD3965 and VB124 (**Fig. S3B**). These findings align with previous studies indicating compensatory roles for MCT1 and MCT4 in lactate export (*39*).

**Fig. 3.**
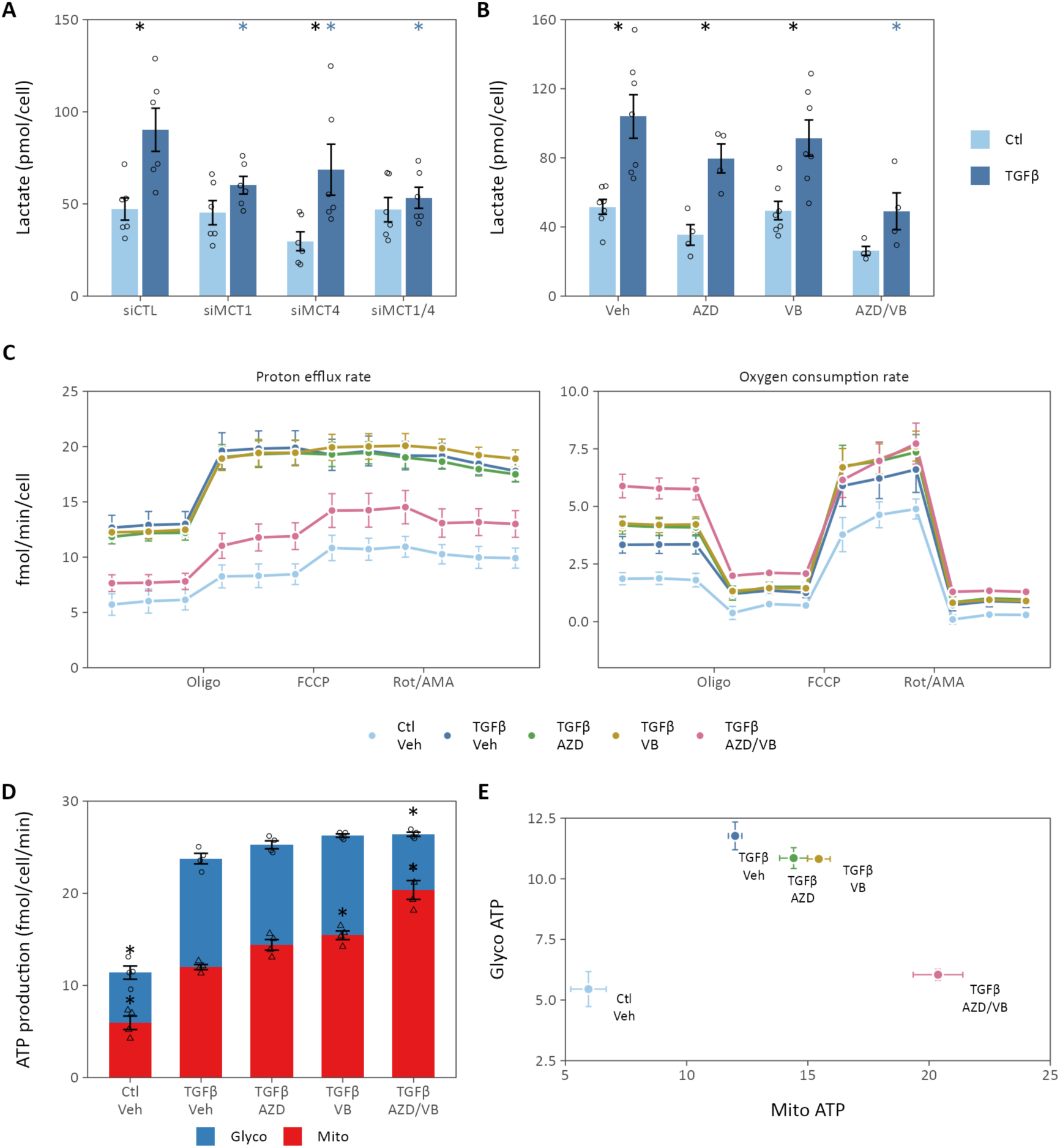
Lactate transport inhibition alters cellular bioenergetics. (**A**) Extracellular lactate was determined by enzymatic assay following TGFβ stimulation of cells treated with siRNA targeting MCT1 or MCT4, separately or together (N = 6 biological replicates, * adjusted p- value < 0.05, *black* compares TGFβ *v.* Ctl, *colored* compares siMCT *v.* siCTL). (**B**) TGFβ-stimulated lactate secretion was measured following treatment with MCT1 inhibitor AZD3965 (AZD), MCT4 inhibitor VB124 (VB), or both (N = 4-13 biological replicates, * adjusted p-value < 0.05, *black* compared TGFβ *v.* Ctl, *colored* compares Drug *v.* Veh). (**C**) Oxygen consumption (OCR) and proton efflux (PER) rates of lung fibroblasts treated with TGFβ and MCT inhibitors for 48 h prior to the assay. Measurements were performed at baseline and following injection of ATP synthase inhibitor oligomycin (Oligo), mitochondrial membrane uncoupler FCCP, and Complex I and III inhibitors rotenone and antimycin A (Rot/AMA) (N = 4 biological replicates, data are mean ± SEM). (**D**) Glycolytic (Glyco) and mitochondrial (Mito) ATP production rates were calculated from PER and OCR, respectively (* adjusted p-value < 0.05 compared to TGFβ/Veh). (**E**) Energy phenogram derived from data presented in (D). TGFβ increases ATP production from both glycolysis and oxidative phosphorylation, favoring the former, while MCT inhibition increases mitochondrial ATP production. Summary data are mean ± SEM.

Furthermore, these results suggest that glycolysis inhibition is not the principal mechanism underlying the attenuation of myofibroblast differentiation by lactate transport inhibition.

To further elucidate the metabolic consequences of lactate transport inhibition, we next measured proton efflux (PER) and oxygen consumption (OCR) rates in lung fibroblasts treated with TGFβ in combination with MCT inhibitors (**Fig. 3C, S3C**). Given that MCTs co-transport protons with lactate, PER serves as a surrogate measure of lactate efflux. In line with previous findings (*8–10, 16, 40*), we observed increases in both PER and OCR following 48 h of TGFβ stimulation, indicative of increased glycolysis and oxidative phosphorylation. Consistent with our direct measures of extracellular lactate and glucose, concurrent treatment with AZD3965 and VB124 was necessary to reduce PER. Consequently, this reduction in PER coincided with an increase in OCR as myofibroblasts transitioned their metabolism from glycolysis to oxidative phosphorylation.

Interestingly, individual administration of AZD3965 and VB124 unexpectedly increased OCR without corresponding decreases in PER. Indeed, the primary consequence of MCT1 or MCT4 inhibition alone was increased cellular ATP production rates driven by upregulation of oxidative phosphorylation (**Fig. 3D-E**). MCT inhibition decreased spare respiratory capacity, indicating that the basal respiratory rate of treated cells approached their maximal oxidative capacity (**Fig. S3D**). No significant differences were observed in glycolytic capacity or electron transport chain coupling efficiency (**Fig. S3D**). Together, these data suggest that the principal metabolic effect of MCT inhibition is the stimulation of oxidative phosphorylation rather than inhibition of glycolysis.

To further test this hypothesis, we performed liquid chromatography-mass spectrometry-based profiling of extracellular and intracellular metabolites from cells treated with AZD3965 and VB124. In accordance with the Seahorse analysis, inhibiting a single lactate transporter had modest effects on extracellular metabolite levels (**Fig. 4A-D, S4**). This analysis confirmed the results of extracellular lactate measurements by enzyme assay, which showed that dual inhibition was required to decrease lactate efflux (**Fig. 4B**). Besides lactate, dual inhibition of lactate transporters primarily altered transcellular fluxes of amino acids (**Fig. 4C-D**), including several metabolites that were differentially regulated by TGFβ treatment (**Fig. S4A**) and dual lactate transport inhibition, including leucine, alanine, ornithine, and ketoleucine.

**Fig. 4.**
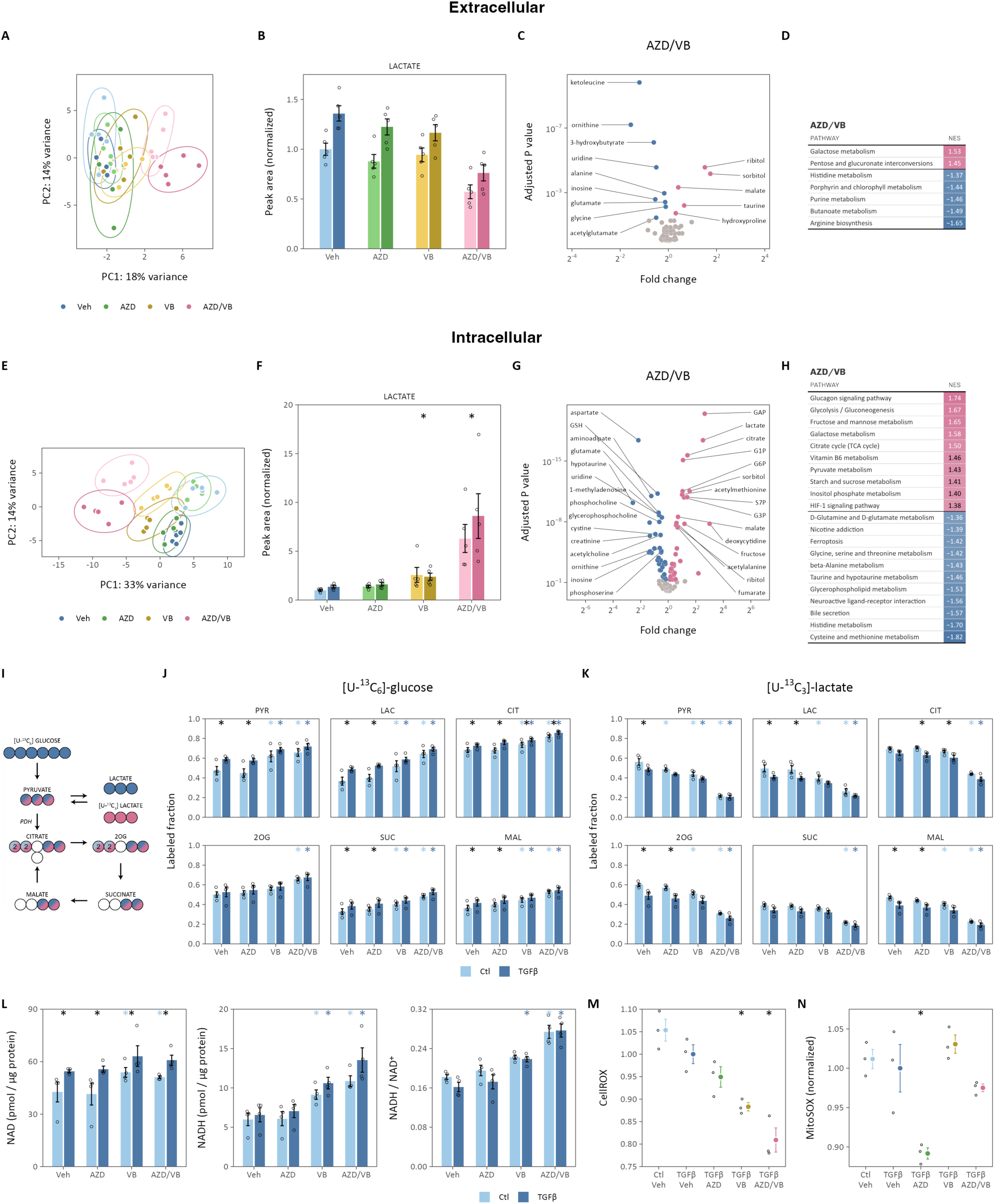
Lactate transporter inhibition promotes oxidative phosphorylation. (**A**) Principal components analysis of metabolites in conditioned medium from lung fibroblasts treated with TGFβ (*bold colors*) or vehicle (*light colors*) and lactate transport inhibitors (N = 5). (**B**) Extracellular lactate determined by LC-MS. (**C**) Volcano plot of significantly altered extracellular metabolites following combined treatment with AZD3965 and VB124. Differentially regulated metabolites are colored (adjusted p-value < 0.1), the top 10 up- and down- regulated of which are labeled. (**D**) KEGG pathways significantly enriched (adjusted p-value < 0.1) with metabolites differentially regulated by combined MCT1 and MCT4 inhibition ordered by normalized enrichment score (NES). Positive NES indicates enrichment in AZD/VB-treated cells while negative NES indicates enrichment in Vehicle-treated cells. (**E**) Principal components analysis of intracellular metabolites extracted from lung fibroblasts stimulated with TGFβ (*bold colors*) or vehicle (*light colors*) in the presence of lactate transport inhibitors (N = 5). (**F**) Intracellular lactate increases significantly with MCT4 inhibition alone (VB) or when combined with MCT1 inhibition (AZD/VB) (* adjusted p-value < 0.05 compared to vehicle control for the main effect of the inhibitor). (**G**) Volcano plot of significantly altered metabolites following combined treatment with AZD3965 and VB124. Differentially regulated metabolites are colored (adjusted p-value < 0.1), the top 10 up- and down-regulated of which are labeled. (**H**) KEGG pathways significantly enriched (adjusted p-value < 0.1) with metabolites differentially regulated by combined MCT1 and MCT4 inhibition ordered by NES. Positive NES indicates enrichment in AZD/VB-treated cells while negative NES indicates enrichment in Vehicle-treated cells. (**I**) Schematic of ^13^C isotope labeling of carbon atoms from glucose (*blue*) or lactate (*pink*) into tricarboxylic acid cycle metabolites. Each circle represents a carbon atom. (**J**-**K**) Stable isotope incorporation from [U-^13^C_6_]-glucose (J) or [U-^13^C_3_]-lactate (K) into the intracellular metabolites pyruvate (PYR), lactate (LAC), citrate (CIT), 2-oxoglutarate (2OG), succinate (SUC), and malate (MAL) (N = 4 biological replicates, * adjusted p-value < 0.05, *black* compares TGFβ *v.* Ctl for a given treatment, *colored* compares treatment *v.* vehicle for the indicated condition). (**L**) Intracellular lactate oxidation is coupled to NADH/NAD^+^, which was determined by enzymatic cycling assay (N = 4 biological replicates; * adjusted p-value < 0.05; *black* compares TGFβ to control within a given treatment, *colored* compares the treatment effect to control for a given condition). (**M**) Normalized CellROX fluorescence, an indicator of cellular reactive oxygen species, normalized to TGFβ-treated cells (N = 3 biological replicates; * adjusted p-value < 0.05 compared to TGFβ-treated cells). (**N**) MitoSOX fluorescence, a marker of mitochondrial superoxide production, was measured and normalized to MitoTracker fluorescence, a marker of mitochondrial mass (N = 3 biological replicates; * adjusted p-value < 0.05 compared to TGFβ-treated cells). Summary data are mean ± SEM.

Similar effects were observed on intracellular metabolites. PCA showed distinct clustering of treatment groups, with the drug effects aligning with PC1 and TGFβ treatment effects with PC2 (**Fig. 4E**). Similar to the extracellular flux results, the magnitude of drug-induced perturbations increased from MCT1 to MCT4 to combined inhibition (**Fig. 4E**), reflected by intracellular lactate levels (**Fig. 4F**). As expected from extracellular lactate measures following MCT inhibition, intracellular lactate accumulated moderately with MCT4 inhibition and substantially with MCT1/4 inhibition. TGFβ treatment resulted in diverse changes in the intracellular metabolomic profile of treated fibroblasts (**Fig. S4C-D**). The effect of AZD3965 alone on intracellular metabolite levels was modest (**Fig. S4G-H**). Conversely, MCT4 inhibition by VB124 alone (**Fig. SK-L**) or in combination with AZD3965 (**Fig. 4G-H**) caused substantial perturbations to intracellular metabolism. Specifically, we observed enrichment of the glycolysis and tricarboxylic acid (TCA) cycle metabolite sets with MCT4 inhibition. These findings collectively suggest that inhibiting lactate export leads to the accumulation of upstream glycolytic intermediates that are rerouted to mitochondrial oxidative metabolic pathways.

To test this hypothesis, we labeled lung fibroblasts with [U-^13^C_6_]-glucose (8 mM) in medium containing lactate (2 mM), glutamine (1 mM), and pyruvate (1 mM) during TGFβ stimulation and treatment with MCT inhibitors (**Fig. 4J, S5**). TGFβ increased ^13^C incorporation from glucose into pyruvate, lactate, citrate, succinate (SUC), and malate (MAL), indicating enhanced flux from glucose into the TCA cycle. While AZD3965 had minimal impact on these labeling patterns, MCT4 inhibition significantly elevated the fractions of these metabolites labeled by ^13^C, providing direct evidence for a proportional increase in glucose oxidation following MCT4 inhibition, consistent with the results of our bioenergetic and steady-state metabolomics experiments described above.

Increased metabolite labeling from [U-^13^C_6_]-glucose must be offset by decreased labeling from other substrates. Given recent data suggesting lactate as a major oxidative fuel source in the lung (*41–43*), we hypothesized that MCT inhibition would decrease exogenous lactate oxidation. To test this, lung fibroblasts were cultured with [U- ^13^C_3_]-lactate (2 mM) in medium containing naturally labeled glucose, glutamine, and pyruvate (**Fig. 4K, S5**).

Extracellular [U-^13^C_3_]-lactate labeled approximately 50% of intracellular pyruvate and lactate at baseline, with significant downstream incorporation into TCA metabolites. This labeling decreased following TGFβ treatment, mirroring increased fractional labeling from glucose (**Fig. 4J**). MCT inhibition had no impact on fractional labeling of TCA intermediates by [U-^13^C_5_]-glutamine (**Fig. S4M, S5**). MCT4 inhibition alone or in combination with AZD3965 decreased ^13^C labeling of intracellular metabolites by lactate. These findings demonstrate the importance of MCT4 for lactate import at physiologic lactate concentrations, in contrast to the prevailing sentiment that MCT4 is primarily a lactate exporter. Moreover, these data further underscore the relatively greater importance of MCT4 activity in fibroblast metabolism compared to MCT1. Consistent with our measurements of extracellular lactate, the effects of MCT inhibition were more pronounced when both inhibitors were used simultaneously, again highlighting some functional redundancy of MCT1 and MCT4 in these cells.

### Lactate transport inhibition contributes to antioxidant defense mechanisms

Lactate metabolism is closely coupled to cellular redox homeostasis through its metabolism by lactate dehydrogenases, which transfer electrons from lactate to NADH. Building on previous findings highlighting the significance of ROS in TGFβ-mediated gene expression (*44, 45*), we investigated the impact of lactate transport inhibition on cellular redox balance. Consistent with increased intracellular lactate upon MCT4 inhibition, we observed a corresponding rise in intracellular NADH/NAD^+^ (**Fig. 4L**), coupled with a reduction in total ROS as measured using the CellROX fluorescent probe (**Fig. 4M**). AZD3965, but not VB124, decreased mitochondrial superoxide production (**Fig. 4N**). Neither inhibitor affected mitochondrial biomass measured by MitoTracker fluorescence (**Fig. S6B**). Less substantial changes were observed in the NADPH/NADP^+^ ratio, where VB124 decreased NADPH/NADP^+^ in TGFβ-treated cells (**Fig. S6A**). Notably, TGFβ did not induce ROS production in our experimental system.

Previous research has proposed proline biosynthesis as a mechanism for NIH-3T3 fibroblasts to mitigate ROS accumulation following TGFβ stimulation (*45*). In this model, proline synthesis from glutamine consumes reducing equivalents from NADPH and NADH, thereby ameliorating reductive stress and decreasing ROS production. Contrary to these findings, we did not observe TGFβ-induced proline elevations in primary lung fibroblasts (**Fig. S6C**). Overall, the fractional labeling of proline from [U-^13^C5]-glutamine was modest at 10% compared to the previously reported 40% (**Fig. S6D**). While we noted trends toward increased proline production from glutamine with MCT4 inhibition, this mechanism appears insufficient to explain the effects of MCT inhibitors on ROS generation.

### Lactate transport inhibition does not alter classical TGFβ signaling pathways

Next, we examined TGFβ-dependent signaling, which activates both SMAD and non-SMAD signaling pathways through a cascade of protein phosphorylation events. Following 48 h of TGFβ stimulation, MCT inhibitors did not diminish Smad3 or ERK phosphorylation (**Fig. 5A-B**).

**Fig. 5.**
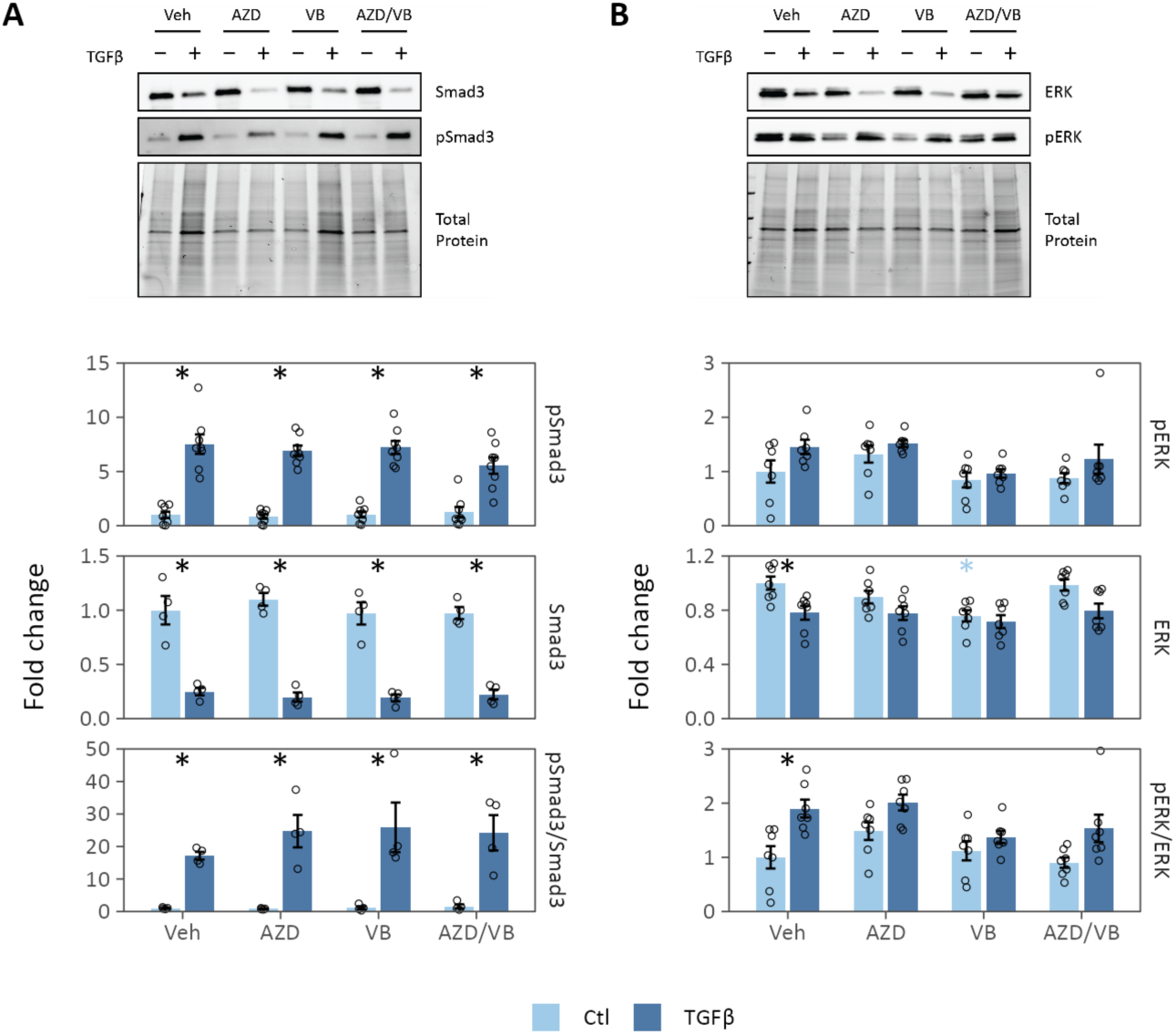
Lactate transport inhibition does not alter TGFβ signaling pathways. (**A**) Phosphorylation of canonical TGFβ receptor target, Smad3, in lung fibroblasts treated with MCT inhibitors. (**B**) Phosphorylation of non-canonical TGFβ receptor target, ERK, in lung fibroblasts treated with MCT inhibitors. Summary data are mean ± SEM (* adjusted p-value < 0.05; *black* compares TGFβ to control within a given treatment, *colored* compares the treatment effect to control for a given condition).

Given that increased intracellular lactate correlated with more potent inhibition of α-SMA expression, we treated cells with TGFβ in combination with extracellular lactate (10 mM). A previous study suggested that extracellular lactate modestly increases α-SMA expression independently of TGFβ (*13*). However, we observed no impact of extracellular lactate on α-SMA expression in TGFβ-treated lung fibroblasts (**Fig. S7A**), indicating intracellular lactate accumulation alone does not mediate the antifibrotic effects of MCT inhibition.

Prior research has linked inhibition of glycolysis and lactate production to hypoxia-inducible factor 1α activation (HIF-1α), contributing to myofibroblast differentiation (*10, 13*). We observed maximal HIF-1α activation 6 h after TGFβ stimulation. At this time point, HIF-1α protein levels remained consistent across vehicle-, AZD3965-, and VB124-treated cells (**Fig. S7B**). However, in lung fibroblasts treated with dual inhibitors, HIF-1α protein increased compared to vehicle, consistent with the enrichment of the “Hypoxia” gene set in our RNA-seq analysis (**Fig. 2F**). These findings, alongside previous studies, suggest that lactate transport inhibition acts downstream of HIF-1α- dependent transcriptional programs to inhibit myofibroblast differentiation.

### MCT inhibition decreases experimental pulmonary fibrosis

Building on our *in vitro* findings indicating an antifibrotic effect of lactate transporter inhibition, we proceeded to evaluate the efficacy of MCT inhibitors in a bleomycin-induced mouse model of pulmonary fibrosis. Mice received 1.2 U/kg bleomycin by intratracheal administration. Seven days later, the animals began treatment with AZD3965 (100 mg/kg twice daily) or VB124 (30 mg/kg once daily) or vehicle by oral gavage (**Fig. 6A**).

**Fig. 6.**
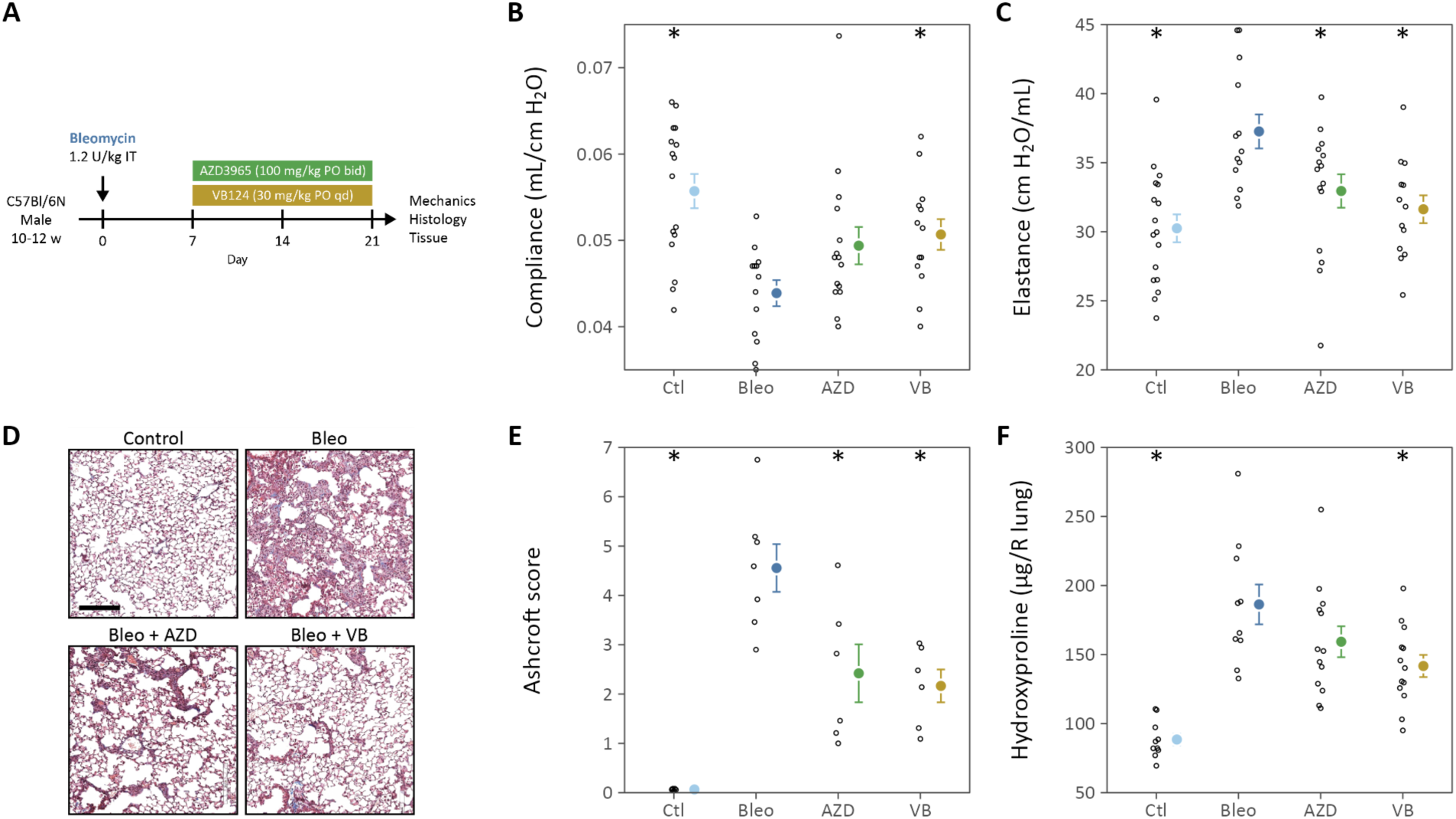
Lactate transporter inhibition attenuates bleomycin-induced pulmonary fibrosis. (**A**) C57Bl/6N mice were administered bleomycin (1.2 U/kg) on day 0. Beginning on day 7, animals were treated with AZD3965 (AZD) or VB124 (VB) for 14 days prior to euthanasia on day 21. On day 21, mice were anesthetized with pentobarbital for lung function measurements. (**B**) Static compliance of the lung was measured by pressure-volume loop analysis. (**C**) Elastance was measured using the forced oscillation technique for 8 s (Prime-8) (N = 13-17). (**D**) Representative trichrome-stained histologic sections of mouse lungs. Scale bar is 200 μm. (**E**) Ashcroft scores from histologic sections from (D) determined by a lung pathologist blinded to treatment assignment (N = 6-7). (**F**) Right lung hydroxyproline content (N = 10-13). Data points show individual mice, summary statistics show the mean ± SEM, * adjusted p-value < 0.05 compared to bleomycin-treated (Bleo) vehicle control.

Considering likely toxicity, we did not assess the combination of inhibitors. Compared to vehicle, mice treated with VB124 had increased weight recovery 21 days after bleomycin administration (**Fig. S8**). Lung mechanics improved by approximately 50% of baseline compared to vehicle controls following 14 days of MCT inhibitor treatment (**Fig. 6B-C**). This improvement in measures of lung stiffness was further supported by histologic assessment of fibrosis severity using Ashcroft scoring (*46*) (**Fig. 6D-E**) and hydroxyproline content measurement (**Fig. 6F**). Taken together, these data demonstrate substantially decreased pulmonary fibrosis severity after 14 days of treatment with lactate transport inhibitors.

### MCT inhibition reprograms lung metabolism *in vivo*

To explore metabolic changes following lactate transporter inhibition *in vivo*, we conducted metabolomic profiling of lung and plasma from mice treated with bleomycin and MCT inhibitors (**Fig. S9**). While bleomycin administration substantially altered the lung metabolic profile, treatment with AZD3965 or VB124 had little additional impact on total metabolite levels. However, unlike bleomycin alone or following AZD3965 treatment, VB124 treatment resulted in significant alterations in several circulating metabolites (**Fig. S9I**). Among these, VB124 significantly increased circulating lactate, while AZD3965 led to decreased circulating lactate (**Fig. 7A**).

**Fig. 7.**
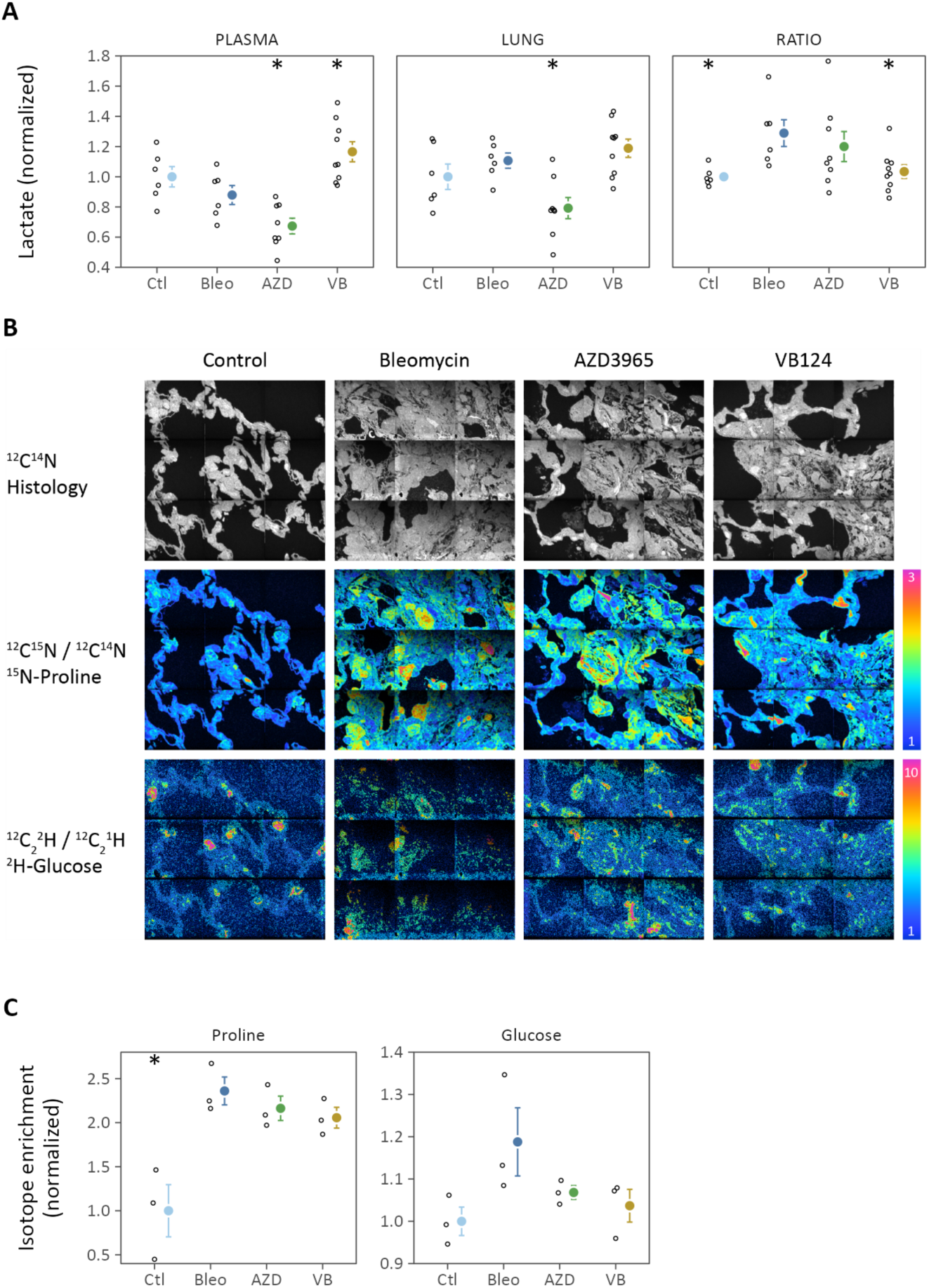
Lactate transporter inhibition reprograms lung metabolism *in vivo*. (**A**) Plasma and lung lactate levels following 14-day treatment with AZD3965 or VB124 (N = 6-9; * adjusted p-value < 0.05 compared to bleomycin control). (**B**) Representative images from multi-isotope imaging mass spectrometry (MIMS). The ^12^C^14^N ion illustrates tissue histology. The ^12^C^15^N/^12^C^14^N ion ratio image shows enrichment from ^15^N-proline (colorbar 1-3-fold of natural abundance). The ^12^C2^2^H/^12^C2^1^H ratio shows enrichment above natural abundance from ^2^H-glucose (colorbar 1-10-fold of natural abundance). (**C**) Quantification of isotope enrichment (N = 3 biological replicates, * adjusted p-value < 0.05 compared to bleomycin-treated control).

This trend was also reflected in the lung, where only modest changes were noted following bleomycin, contrary to prior reports (*13*). Considering the influence of our treatments on circulating lactate, we calculated the lung- to-plasma lactate ratio. This analysis revealed the anticipated increase in lactate in fibrotic lungs, with levels returning to baseline following the addition of VB124 (**Fig. 7A**).

We were surprised to observe relatively few metabolic changes in whole lung lysates following MCT inhibition compared to our *in vitro* findings. We speculated that this discrepancy might arise from measuring steady-state metabolite levels that do not reflect differences in metabolic flux. To test this hypothesis, we performed multi- isotope imaging mass spectrometry (MIMS) of mouse lungs following administration of ^15^N-proline and either ^2^H-glucose or ^13^C-glucose. MIMS enables the quantification of stable isotope tracer flux into tissue biomass with subcellular spatial resolution (*47, 48*). For three days preceding tissue collection, mice received twice daily intraperitoneal injections of 5 mg ^15^N-proline as a fibrosis tracer and 50 mg glucose isotope as a metabolic tracer. Subsequently, lung tissue sections were imaged by nanoscale secondary ion mass spectrometry to quantify spatially resolved isotope tracer uptake (**Fig. 7B-C**). This labeling approach proved effective, demonstrating approximately 1.5-fold enrichment ^15^N, 4.6-fold for ^2^H, and 2.4-fold for ^13^C above natural isotope abundance. Pulmonary fibrosis correlated with a significant increase in ^15^N labeling from proline, consistent with increased collagen synthesis and deposition during the labeling period. Similarly, drug-treated animals exhibited less ^15^N incorporation per tissue area, consistent with lung function and histological analyses. Glucose labeling displayed a similar pattern, wherein lactate transporter inhibition led to reduced glucose incorporation into tissue biomass. Importantly, these findings are not merely attributable to decreased tissue fibrosis, as we selectively imaged more fibrotic areas (**Fig. 7B**) and normalized the enrichment values to tissue area.

### VB253, a novel MCT4 inhibitor, alleviates experimental pulmonary fibrosis

Our preclinical findings indicate that MCT4 inhibition, as a single therapeutic target, exhibits greater antifibrotic efficacy than MCT1 inhibition, both in suppressing myofibroblast differentiation *in vitro* and reducing bleomycin- induced fibrosis. VB253 is a novel inhibitor of MCT4 with ∼10-fold increased selectivity for MCT4 *v*. MCT1 and ∼10-fold increased potency for MCT4 inhibition. Like VB124, VB253 dose-dependently decreased TGFβ- stimulated α-SMA in human IPF lung fibroblasts *ex vivo* (**Fig. 8A**). To assess its efficacy compared to established therapies, we compared VB253 to nintedanib, a clinically approved antifibrotic medication for pulmonary fibrosis (**Fig. 8A-B**). Nintedanib inhibits TGFβ-mediated myofibroblast differentiation and decreases Col1a1 expression *in vitro* and *in vivo* (*49*). While both compounds effectively attenuated α-SMA expression, nintedanib exhibited moderate cytotoxicity absent with VB253 (**Fig. 8B**). Nintedanib mitigates myofibroblast differentiation partly by inhibiting TGFβ receptor phosphorylation and Smad-dependent signaling pathways (*50*).

**Fig. 8.**
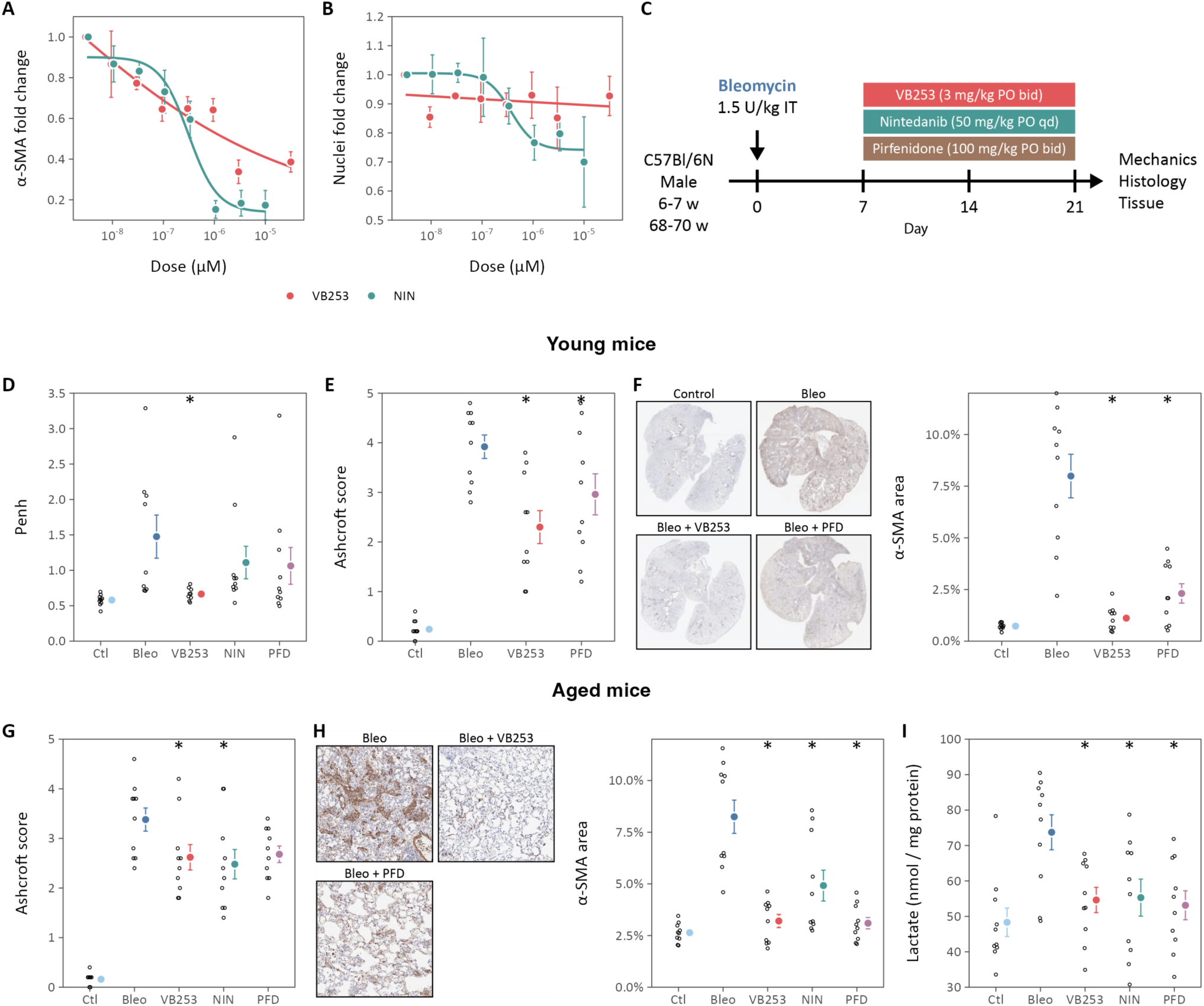
VB253, a novel MCT4 inhibitor, is potently antifibrotic *in vitro* and *in vivo*. (**A**) Dose-response of α-SMA protein expression in IPF lung fibroblasts treated with VB253 or nintedanib (NIN). (**B**) Number of nuclei, a marker of cell viability, following treatment with VB253 or NIN. Summary data are mean ± SEM of cell responses from N = 3 different donors. (**C**) Young (8-10 w) or aged (60+ w) mice were treated with bleomycin (1 U/kg) on day 0. Beginning on day 7, animals were treated with VB253 or the FDA-approved antifibrotics nintedanib (NIN) and pirfenidone (PFD) for 14 days prior to euthanasia on day 21. (**D**) Enhanced pause (Penh) is a non-invasive index of mouse tidal breathing that increases with bleomycin treatment. (**E**-**G**) Histologic assessment of VB253-treated mouse lungs demonstrates decreased fibrosis by Ashcroft score (E) and decreased α-SMA protein expression (F). (**G**-**I**) Similar histologic improved was observed in aged mice with decreased histologic fibrosis (G) and decreased α-SMA protein expression (H). (**I**) VB253 decreased lung lactate, similar to established antifibrotics. Data points show individual mice, summary statistics show the mean ± SEM, * adjusted p-value < 0.05 compared to bleomycin-treated (Bleo) vehicle control.

Correspondingly, nintedanib dose-dependently attenuated TGFβ-mediated Smad3 phosphorylation, while VB253 and VB124 had no impact on this signaling pathway (**Fig. 5, S10**).

Subsequently, we evaluated the potential of VB253 to counter bleomycin-induced pulmonary fibrosis. In experiments conducted independently from those reported above, VB253 was administered to young mice (8-10 weeks) beginning on day 7 following bleomycin (**Fig. 8C**). The existing pulmonary fibrosis therapies, nintedanib and pirfenidone, served as comparators. Three weeks after bleomycin, whole-body plethysmography was performed to assess breathing patterns in unrestrained mice. Enhanced pause (Penh) is a dimensionless index describing airflow during tidal breathing found to increase significantly in bleomycin-treated mice (*51*).

VB253 restored Penh to baseline levels, suggesting normalization of respiratory patterns in bleomycin-treated mice (**Fig. 8D**). Histologic assessment confirmed reduced pulmonary fibrosis (**Fig. 8E**) and α-SMA expression (**Fig. 8F**). Moreover, the effects of VB253 were comparable to the antifibrotic effects of nintedanib and pirfenidone with decreased cytotoxicity in *in vitro* assays.

Compared to young mice, aged mice exhibit more severe and persistent bleomycin-induced pulmonary fibrosis (*52*). Given the clinical relevance of age-related IPF incidence, we investigated the antifibrotic effects of VB253 in aged (60+ weeks) mice. Similar to young mice, VB253 decreased fibrosis severity, as quantified by Ashcroft score and α-SMA expression (**Fig. 8G-I**). The magnitude of improvement paralleled that of nintedanib and pirfenidone. As expected, VB253 decreased total lung lactate akin to VB124 (**Fig. 7A**). Collectively, these findings provide compelling preclinical evidence supporting lactate transporter inhibition as a novel therapeutic strategy for fibrotic lung disease.

## DISCUSSION

Our findings identify the pivotal role of lactate transport in the metabolic reprogramming associated with myofibroblast differentiation both *in vitro* and *in vivo*. Elevated expression of the lactate transporters MCT1 and MCT4 was observed in IPF lung explants and experimental models, underscoring their significance. Inhibiting these transporters mitigated TGFβ-stimulated myofibroblast differentiation and attenuated the severity of bleomycin-induced pulmonary fibrosis without a demonstrable impact on classical TGFβ receptor signaling pathways. Metabolically, MCT antagonists promoted glucose oxidation while reducing glucose carbon incorporation into fibrotic lung regions, correlating with decreased oxidative stress. MCT4 inhibition consistently exhibited superior antifibrotic potency compared to MCT1 inhibition. We introduce a novel MCT4 inhibitor, VB253, which has a more favorable pharmacologic profile than VB124, and is currently undergoing Phase 1 clinical trials. Altogether, these data establish lactate transport as a promising metabolic target for therapeutic intervention in pulmonary fibrosis.

Metabolic reprogramming characterizes myofibroblast differentiation, with previous studies revealing alterations in carbohydrate, amino acid, and lipid metabolic pathways that promote fibrogenesis (*7*). Of these, increased lactate production emerged as an early metabolic hallmark associated with pulmonary fibrosis (*13*). Subsequent investigations identified glycolysis activation as the driving force behind increased lactate production by myofibroblasts (*9*). Furthermore, inhibition of glycolysis not only prevented myofibroblast differentiation (*8, 9, 14*), but also attenuated experimental pulmonary fibrosis (*15, 16*). Our findings align with this metabolic shift, as we observed upregulated expression of lactate transporters MCT1 and MCT4, supporting this glycolytic phenotype in myofibroblasts.

MCTs are proton-coupled monocarboxylate symporters with varying affinities for lactate, pyruvate, and other monocarboxylates (*35*). MCT1 is constitutively and ubiquitously expressed and primarily considered to be a lactate importer with a lactate affinity (*K*_M_) ranging from 3-6 mM. By contrast, MCT4 expression is dynamically regulated, including by HIF-1α, and traditionally considered a lactate exporter with a lower lactate affinity (*K*_M_ 30-40 mM). However, recent evidence suggests that MCT4 has a much higher affinity for lactate import than previously appreciated (*K*M 1 mM) (*38*). These findings align with our results where inhibiting both MCT1 and MCT4 was required to block lactate export (**Fig. 3, S3**) and isotope incorporation from extracellular [U-^13^C_3_]- lactate import (**Fig. 4K**). Previous studies in cancer cells have also noted that MCT4-expressing cells are resistant to the cytotoxic effects of MCT1 inhibition (*28*). Interestingly, inhibiting either MCT1 or MCT4 decreased myofibroblast differentiation without affecting lactate export, adding further complexity to their roles in cellular metabolism.

To clarify the metabolic consequences of lactate transporter inhibition, we performed a comprehensive metabolic analysis of myofibroblasts treated with MCT inhibitors, encompassing bioenergetic measurements, metabolomic profiling, and stable isotope tracing. The primary metabolic consequence of MCT inhibition is stimulation of oxidative phosphorylation. MCT inhibitors increased the fraction of mitochondrial ATP production, which was associated with increased levels of TCA intermediates and increased isotope incorporation from glucose. Collectively, these data suggest that MCT inhibition redirects glucose carbon flux away from lactate fermentation and toward glucose oxidation. Several mechanisms could underlie this shift. Given that the observed effects generally scale with intracellular lactate levels, mass action likely plays an important role in driving this metabolic phenomenon. Additionally, lactate accumulation is closely coupled in NADH production through lactate dehydrogenase activity. Cytoplasmic NADH may be transported into the mitochondria through the malate-aspartate shuttle and oxidized by the electron transport chain. Recently, lactate itself was shown activate the mitochondrial electron transport chain independent of its metabolism (*53*), though the molecular mechanism remains unknown.

Our data reveal that lactate transport inhibition exploits novel antifibrotic mechanisms distinct from TGFβ signaling pathways and suggest that these metabolic effects may signal to antifibrotic transcriptional programs by dampening ROS levels. Increased ROS production has been observed following TGFβ treatment and seems to be essential for myofibroblast differentiation (*44, 45*). While our cells and model system did not replicate TGFβ- dependent increases in ROS, we observed decreased CellROX oxidation with MCT4 inhibition and decreased MitoSOX oxidation with AZD3965. Additional signaling mechanisms may also contribute, such as post- translational modification of protein lysines by lactate (*i.e.*, protein lactylation). Histone lactylation serves as an epigenetic modification that stimulates gene expression (*54*) and this modification has been increasingly identified as a critical regulator of protein function, including cytoskeletal proteins (*55*). Furthermore, ongoing research is uncovering novel lactate targets and mechanisms as significant mediators of metabolic signaling (*56, 57*).

Through our comprehensive metabolic investigation of lactate transport inhibition, we have also generated valuable data on the metabolic and transcriptional consequences of human lung myofibroblast differentiation. TGFβ stimulation induced notable alterations in amino acid and nucleic acid metabolic pathways (**Fig. S4**). While previous studies have underscored the significance of glutamine, proline, and taurine (*7, 58, 59*), the role of branched chain amino acid metabolism, for instance, remains unexplored. Integrating these multi-omics data sets could unveil novel molecular targets for future drug development.

In addition to metabolomic profiling, we conducted stable isotope tracing using glucose, lactate, and glutamine substrates in TGFβ-treated primary human lung fibroblasts. To our knowledge, this represents the first comprehensive dataset on intracellular substrate metabolism in this widely employed model of myofibroblast differentiation. Overall, isotope labeling patterns changed little following TGFβ treatment (**Fig. S5**). This observation, coupled with the results of our extracellular flux experiments, suggests that TGFβ primarily enhances metabolite flow through metabolic pathways without substantially altering the pathways themselves. Consistent with increased glucose uptake and accumulation of glycolytic metabolites, a greater fraction of pyruvate, lactate, alanine, and serine are labeled by [U-^13^C6]-glucose, countered by a reduction in the fractional labeling from [U-^13^C3]-lactate. Consistent with our prior findings (*43*), nearly 50% of TCA metabolites (citrate, 2- oxoglutarate, succinate, and malate) are labeled by [U-^13^C3]-lactate, highlighting the importance of lactate as a respiratory fuel source in these cells.

Recently, the contributions of fibroblast metabolic pathways to extracellular matrix production have garnered significant attention (*7*). Serine and glycine synthesis from the glycolytic intermediate 3-phosphoglycerate, and proline synthesis from glutamine, have been implicated in myofibroblast differentiation and pulmonary fibrosis (*11, 12, 45, 58, 60*). Intriguingly, our data indicate limited incorporation of glucose or lactate carbon into serine and minimal glutamine carbon incorporation into proline. These differences may stem from differences in the cell types (IMR-90 fetal lung fibroblasts or NIH-3T3 spontaneously immortalized mouse embryonic fibroblasts) or the culture medium composition (Eagle’s minimum essential medium, which lacks serine, glycine, and proline). Our study suggests that fibroblasts preferentially utilize available amino acids over rerouting substrates into biosynthetic pathways. Developing metabolic flux models using human physiologic medium (*61*) could provide a more accurate and comprehensive understanding of substrate flow into energetic and biosynthetic pathways, better mimicking fibroblast metabolism *in vivo*.

Even more informative than *in vitro* model systems are approaches that enable the study of cell metabolism *in vivo*. Here, we performed metabolomic profiling of lung and plasma samples from mice treated with bleomycin and MCT inhibitors (**Fig. S9**). These analyses did not reveal changes in whole lung metabolite levels, demonstrating a potential limitation of using bulk metabolomics to monitor metabolic changes in heterogeneous cell populations *in vivo*. To overcome this challenge, we employed spatial metabolic imaging.

Mice were administered stable isotope tracers of proline and glucose before euthanasia. Since these tracers are administered to live animals, multi-isotope imaging mass spectrometry identifies lung regions that are metabolically active during the labeling period. Since tissues were fixed and processed *ex vivo*, the isotope signals indicate substrate incorporation into fixable biomass (*62, 63*). Thus, using multi-isotope imaging mass spectrometry, we directly quantify isotope flux from glucose into fibrotic lung regions.

Our findings demonstrated that ^15^N-proline enrichment levels correlated with tissue fibrosis as assessed histologic and biochemically. This finding emphasizes the ability of cells to utilize circulating proline for protein synthesis, potentially diminishing the significance of *de novo* proline biosynthesis *in vivo*. Moreover, the ^2^H- glucose signal provided direct evidence for MCT-dependent metabolic reprogramming in mice, showing reduced carbohydrate incorporation into matrix proteins. Owing to the financial and time costs of isotope tracing and imaging, we were able to study only a few animals per group. Nevertheless, metabolic imaging at high spatial resolution holds great promise for correlating cell identities from spatial transcriptomic profiles with metabolic features. Furthermore, stable isotopes may be safely administered to human patients prior lung biopsy or explant (*47, 64*), offering a strategy that could significantly advance our understanding of how cell metabolism contributes to pulmonary fibrosis.

MCT1 and MCT4 are expressed by many cells in the lung, notably macrophages and dendritic cells (*65*). Although our *ex vivo* and *in vitro* experiments suggest that inhibiting myofibroblast differentiation is a primary antifibrotic mechanism, the inhibition of lactate transporters expressed by other cell types may also contribute to their therapeutic effect. For example, MCT4 expression is upregulated as part of a HIF-1α gene expression signature in transitional AT2 cells that accumulate in pulmonary fibrosis and contribute to aberrant repair processes (*66, 67*). Future work will explore the cell-type-specific effects of these transporters in conditional knockout mice.

The poor pharmacologic properties of small molecules studied previously have prevented translation into metabolic therapies for human pulmonary fibrosis. Both AZD3965 and VB253 exhibit low IC_50_ values of approximately 2 nM, compared to the next most potent inhibitor studied in pulmonary fibrosis models, lonidamine, with an IC_50_ of 7,000 nM (*68*). As metabolic targets downstream of glycolysis, lactate transport inhibitors also offer better tolerance by allowing glycolysis to continue supporting glucose oxidation, contrasting with upstream glycolysis inhibitors that more severely impact cellular bioenergetics. Pharmacologic interest in lactate transporters has been driven by the recognition of increased lactate transporter expression in a variety of cancers (*69*). AZD3965 was selected for this study as it has been investigated in human clinical trials for advanced solid organ malignancies (*22*). In this Phase 1 study, AZD3965 was generally well-tolerated with 7 of 40 patients experiencing dose-limiting toxicities including asymptomatic, reversible ocular changes; acidosis; and increased troponin. VB124 was the first selective MCT4 inhibitor developed (*21*) and we now introduce VB253 as a second generation MCT4 inhibitor. Our data demonstrate that VB253 has similar efficacy in experimental pulmonary fibrosis models as the established antifibrotics, nintedanib and pirfenidone, with potentially less cytotoxicity (**Fig. 8**). MCT4 global knockout mice are viable and breed normally (*70, 71*), raising optimism that VB253 will be well tolerated in humans, and a Phase 1 clinical trial of VB253 is currently underway.

In summary, our findings highlight the pivotal role of lactate transporter in driving myofibroblast differentiation and pulmonary fibrosis. Through a comprehensive metabolic phenotyping approach, we have characterized the antifibrotic mechanisms associated with MCT inhibition and provided compelling evidence of metabolic reprogramming in animal models. Furthermore, we have validated the antifibrotic effectiveness of existing lactate transport inhibitors using established preclinical disease models. Altogether, our results significantly advance lactate transport inhibition as a promising therapeutic approach for patients suffering from pulmonary fibrosis.

## MATERIALS AND METHODS

### Study Design

This study was designed to investigate the role of lactate transporters in myofibroblast differentiation and pulmonary fibrosis. The objectives of this study were (i) to determine the expression of lactate transporters in IPF lungs and model systems, (ii) to characterize the phenotypic effects of lactate transport inhibition in model systems, and (iii) to profile the metabolic consequences of lactate transport inhibitors.

Cell culture experiments were performed at least three times. The number of animals per experimental group was chosen based on prior publications and experiments were repeated at least once. Stable isotope tracing experiments were performed on three animals per group owing to resource availability. Animals were randomly assigned to treatment. The pathologist scoring histologic fibrosis severity was blinded to treatment assignment. The number of unique patient samples was determined by clinical availability.

### Statistical Analysis

Data analysis, statistical comparisons, and visualization were performed in R (*72*). Experiments included technical and biological replicates as noted in the Materials and Methods. The number of biological replicates (N) is indicated in the figure legends. Summary data show the mean ± SEM. Outliers were identified using twice the median absolute deviation as a cutoff threshold. Comparisons were performed using linear mixed-effects models with condition (±TGFβ), treatment, and their interaction as fixed effects and biological replicate or donor as a random effect. Significant differences in estimated marginal means were identified by comparisons to the multivariate *t* distribution. Metabolomics and RNA-seq data were analyzed as described in the Materials and Methods. Probability values less than 0.05 were considered statistically significant.

### Study approval

Human samples were obtained through the BWH Biorepository for Understanding Inflammatory Lung Disease (BUILD) or the MGH ILD Translational Research Program and their collection was approved by the Mass General Brigham Institutional Review Board (2013P002332, 2016P001890, 2019P003592, 2020P002765). All animal experiments were approved by the Brigham and Women’s Hospital IACUC (2020N000199).

## LIST OF SUPPLEMENTARY MATERIALS

Materials and Methods

Fig. S1-S10

## ACKNOWLEDGEMENTS

The authors thank Louise Trakimas and the HMS Electron Microscopy Facility for MIMS sample preparation and consultation. The content is solely the responsibility of the authors and does not necessarily represent the official views of the National Institutes of Health. VB124 was a gift from Vettore, LLC to the Oldham laboratory.

## Author contributions

- Conceptualization: KMP, MLS, JR, WMO
- Methodology: DRZ, FL, KMP, CG, JV, BAM, MLS, RSK, JR, WMO
- Investigation: DRZ, FL, KMP, NMK, KJL, CG, JV, LPH, RSK, WMO
- Visualization: WMO
- Funding acquisition: KMP, JR, WMO
- Project administration: KMP, JR, WMO
- Resources: DRZ, KMP, NMK, RMB, NJP, EYK, RSK
- Supervision: KMP, JR, WMO
- Writing - original draft: WMO
- Writing - review and editing: DRZ, FL, KMP, NMK, CG, JV, RMB, BAM, NJP, EYK, MLS, RSK, JR, WMO

## Data and materials availability

The raw data and annotated analysis code necessary to reproduce this manuscript are contained in an R package research compendium available from the Oldham Lab GitHub repository (github.com/oldhamlab/Ziehr.2023.ipf.mcti).

RNA sequences were deposited in the NIH SRA (PRJNA1011992). Details of the processing pipeline and summarized data are available from the Oldham Lab GitHub repository (github.com/oldhamlab/rnaseq.lf.tgfb.mcti).

