## Supplementary Material for "Lactate transport inhibition therapeutically reprograms fibroblast metabolism in experimental pulmonary fibrosis"

#### MATERIALS AND METHODS

##### Key Resources

| Resource | Designation | Source | Identifiers | Notes |
|---|---|---|---|---|
| antibody | HRP-anti-Rabbit IgG | Cell Signaling Technologies | 7074 | 1:5000 |
| antibody | HRP-anti-Mouse IgG | Cell Signaling Technologies | 7076 | 1:5000 |
| antibody | anti-MCT1 | Thermo | PA5-72957 | 1:1000 WB |
| antibody | anti-MCT4 | abcam | ab244385 | 1:2000 WB |
| antibody | anti-α-SMA | abcam | ab32575 | 1:2000 WB |
| antibody | anti-Col1a1 | Cell Signaling | 39952S | 1:1000 WB |
| antibody | anti-HIF-1α | BD Biosciences | 610958 | 1:1000 WB |
| antibody | anti-Smad3 | abcam | ab40854 | 1:2000 WB |
| antibody | anti-pSmad3 | abcam | ab52903 | 1:2000 WB |
| antibody | anti-ERK | Cell Signaling Technologies | 4696S | 1:2000 WB |
| antibody | anti-pERK | Cell Signaling Technologies | 4370S | 1:2000 WB |
| siRNA | siMCT1 | Dharmacon | L-007402-00-0005 | |
| siRNA | siMCT4 | Dharmacon | L-005126-02-0005 | |
| siRNA | siCTL | Dharmacon | D-001810-10-05 | |
| chemical | AZD3965 | Medkoo Biosciences | 206040 | |
| chemical | AR-C155858 | Tocris Bioscience | 4960 | |
| chemical | VB124 | Vettore | | |

#### Chemicals

Recombinant human TGFβ was purchased commercially , dissolved in 10 mM citric acid, pH 3.0, filtered, and diluted to 10 µg/mL in PBS with 0.1% BSA prior to aliquoting and storing at -80 °C. AZD3965 (100 nM; Medkoo) and AR-C155858 (200 nM; Tocris) were purchased from commercial vendors. VB124 (10 µM) was a gift from Vettore, LLC. The drugs were prepared in DMSO and used at the indicated doses unless otherwise indicated. These doses were chosen based on prior reports and their target affinities.

#### Human lung tissue and fibroblasts

Human IPF lung samples were obtained at the time of lung transplant. Control samples were prepared from non-fibrotic and unaffected border regions of lung nodule biopsy specimens or from failed donor controls. To isolate human lung fibroblasts, lung tissue was minced, digested with liberase and DNase, and passed through a sterile 70 µm filter. The filtrate was centrifuged at 300 ×*g* and washed once with RPMI medium (Lonza). Cells were plated in DMEM (Lonza) supplemented with 10% FBS, penicillin, and streptomycin. After two passages, medium was changed to FGM-2 (Lonza).

#### Bleomycin model

C57Bl6/N mice aged 10-12 weeks were anesthetized with ketamine (90 mg/kg) and xylazine (5 mg/kg) and administered 1.2 U/kg bleomycin in PBS (Fresenius Kabi) intratracheally. Control animals received an equivalent volume of PBS. For drug treatments, AZD3965 (100 mg/kg twice daily) and VB124 (30 mg/kg daily) were prepared in 0.5% methylcellulose and 0.1% Tween-20 with sonication and administered by oral gavage every day beginning on day 7 following bleomycin. Control animals received vehicle alone twice daily. After 21 d, lung function measurements were performed. Animals were euthanized and plasma samples were obtained by right ventricular puncture and collected in EDTA-coated tubes. The lungs were dissected and perfused with PBS. The right lung was isolated while the left lung was inflated to 20 $cmH_2O$ prior to fixation. The right lungs were snap frozen.

Lung physiology measurements were performed as previously described *(73)*. Briefly, animals were anesthetized with pentobarbital (100 µg/g; Covetrus, Portland, ME) and tracheally cannulated for pulmonary function measurements using a flexiVent (SCIREQ). Animals were ventilated for 3-5 min with 150 breaths per minute at 10 mL/kg and 3 $cmH_2O$ positive end-expiratory pressure. After two total lung capacity maneuvers to prevent atelectasis and standardize volume history, respiratory elastance was measured using two Prime-8 maneuvers;

values reported are the average of these two maneuvers. A pressure-volume loop was then generated, and static compliance was calculated using the Salazar-Knowles equation.

Histologic fibrosis severity was assessed by Ashcroft scoring of trichrome-stained lung sections in a blinded fashion by an experienced lung pathologist *(74)*. The average of twenty 10× fields is reported for each animal.

Lung hydroxyproline content was determined as described previously *(75)*. The four lobes of the R lung were homogenized in PBS with Halt protease inhibitors (Thermo) and hydrolyzed in 6 N HCl at 120 °C for 12 h, with an aliquot added to 1.4% chloramine-T, 10% 1-propanol, and 0.5 M sodium acetate, pH 6.0. After incubating 20 min at room temperature, 1 ml of Ehrlich's solution (1 M p-value-dimethylaminobenzaldehyde in 70% 1-propanol and 20% perchloric acid) was added. Samples were incubated at 65 °C for 15 minutes. Absorbance was measured at 550 nm and hydroxyproline content was interpolated from a standard curve.

**Myofibroblast differentiation *in vitro***

Normal lung fibroblasts were purchased commercially (Lonza). All fibroblasts were cultured in FGM-2 medium (Lonza). Passages 3-8 were used for experiments. Myofibroblast differentiation was induced *in vitro* using recombinant human TGFβ (2 ng/mL) (Peprotech) for 48 h following 24 h serum starvation unless otherwise indicated.

**Isotope labeling**

Cells were seeded and serum starved in MCDB131 medium lacking glucose, glutamine, and phenol red (genDEPOT) which was supplemented with naturally labeled glucose (8 mM), glutamine (1 mM), and lactate (2 mM) ("light" labeling medium). The concentrations of glucose and glutamine match the concentrations of these substrates determined in standard (FGM-2) growth medium. After 24 h, cells were washed with PBS and the medium was changed to "heavy" labeling medium containing [1,2-$^{13}C_2$]-glucose, [U-$^{13}C_6$]-glucose, [U-$^{13}C_5$]-glutamine, or [U-$^{13}C_2$]-lactate (Cambridge Isotope Labs) along with TGFβ and MCT inhibitors for 48 h. Cell lysates were prepared for metabolomics as described below.

**Immunoblotting**

Human and mouse lung samples were mechanically homogenized in RIPA lysis buffer containing Halt Protease Inhibitor Cocktail (Thermo). Cells were washed with one volume of PBS and collected by scraping in PBS. Cell suspensions were centrifuged at 5,000 ×*g* for 5 min at 4 °C. Pellets were lysed in buffer containing Tris 10 mM, pH 7.4, NaCl 150 mM, EDTA 1 mM, EGTA 1 mM, Triton X-100 1% v/v, NP-40 0.5% v/v, and protease inhibitors.

Protein concentrations were determined by BCA Protein Assay (Thermo). Lysates were normalized for protein concentration and subjected to SDS-PAGE separation on stain-free tris-glycine gels (Bio-Rad), cross-linked and imaged with the Chemidoc system (Bio-Rad), transferred to PVDF membranes with the Trans-Blot Turbo transfer system (Bio-Rad), imaged, blocked in 5% blocking buffer (Bio-Rad), blotted in primary and secondary antibodies, and developed using WesternBright ECL (Advansta). Band signal intensity was normalized to total protein per lane as determined from the stain-free gel or membrane images using Image Lab software (Bio-Rad).

**RNA interference**

Lung fibroblasts were reverse transfected with ON-TARGETplus SMARTpool siRNA (Dharmacon) targeting MCT1 or MCT4 using Lipofectamine RNAiMAX (Thermo). A non-targeting siRNA pool was used a transfection control (Dharmacon). For combined siMCT1 and siMCT4 treatment, the total siRNA dose was kept constant. After 24 h, cells were serum starved for an additional 24 h prior to treatment with TGFβ.

**Gel contraction assay**

Lung fibroblasts in FBM and neutralized collagen solution (TeloCol-3, Advanced Biomatrix) were combined 1:2 to yield 100,000 cells/mL and 1.8 mg/mL collagen. This solution was added to 24-well plates (0.5 mL) and allowed to harden for 10 min at 37 °C prior to the addition of 1 mL FBM. Cells were serum starved overnight prior to releasing the gels and treating them with TGFβ and MCT inhibitors in triplicate. Images were acquired after 24 and 48 h using a ChemiDoc imager (Bio-Rad). Gel areas were measured using Image Lab software (Bio-Rad). Data are expressed as the ratio of TGFβ-treated gel area to control gel area.

**Cell count**

Cell counts were estimated from total DNA quantified using the Quant-iT PicoGreen dsDNA Assay Kit (Thermo) as described previously *(4)*.

**Lactate assay**

Lactate was determined using an enzymatic cycling assay resulting in the reduction of resazurin to the fluorescent chemical resorufin. Medium samples were diluted 1:10 in PBS. Samples were combined with a solution of diaphorase (125 mU), lactate dehydrogenase, NAD (250 µM), and resazurin (25 µM) in Tris buffer (100 mM, pH 8.5) then incubated for 30 min at room temperature. Resorufin fluorescence was measured using a SpectraMax i3x microplate reader (Molecular Devices) and lactate concentration was interpolated from a standard curve. Data were normalized to estimated cell count.

**Glucose assay**

Medium samples were diluted 10-fold in PBS. Glucose concentration was determined using the Glucose Colorimetric Assay Kit (Cayman) according to the manufacturer's protocol. Standards were prepared in PBS.

**Seahorse assays**

Lung fibroblasts were seeded in Seahorse assay plates at a density of 20,000 cells per well. On the following day, the cells were serum starved for 24 h prior to treatment with TGFβ and MCT inhibitors for 48 h. On the day of the assay, the medium was exchanged to Seahorse DMEM without phenol red supplemented with 5 mM HEPES, 10 mM glucose, 2 mM glutamine, and 1 mM pyruvate. OCR and extracellular acidification rates (ECAR) were measured at baseline and following the sequential injections of oligomycin (1 µM final), FCCP (2 µM final), and rotenone plus antimycin A (0.5 µM final each) using an XFe24 bioanalyzer (Agilent). At the end of the assay, cells were stained for 30 min with the fluorescent dye Nuclear Green LCS1 (10 µM; Abcam). Stained cells were quantified using a SpectraMax MiniMax 300 Imaging Cytometer (Molecular Devices) and these cell counts were used to normalize the rate data. Proton efflux rate (PER) was calculated from the ECAR and buffer capacity of the medium. PER from glycolysis was determined by subtracting the contribution of oxidative phosphorylation to total PER using the empirically measured $CO_2$ correction factor of 0.41. ATP from glycolysis was assumed to be equal to basal glycolytic PER. ATP from oxidative phosphorylation was assumed to be 2 × 2.75 × oligomycin-sensitive OCR *(76)*.

**Metabolomics**

Metabolomics data acquisition and analysis were performed as described previously *(4)*.

***Metabolite extraction***

Extracellular metabolites were obtained by mixing conditioned medium 1:4 with 100% MeOH precooled to -80 °C. Intracellular metabolites were obtained after washing cells with 2 volumes of ice-cold PBS and floating on liquid nitrogen. Plates were stored at -80 °C until extraction. Metabolites were extracted with 1 mL 80% MeOH pre-cooled to -80 °C. Mouse plasma and lung homogenates were mixed 1:4 with 100% MeOH precooled to -80 °C. Samples were extracted at -80 °C for 4 h. Insoluble material from these samples was removed by centrifugation at 21,000 ×*g* for 15 min at 4 °C. The supernatant was evaporated to dryness at 42 °C using a SpeedVac concentrator (Thermo Savant). Samples were resuspended in 35 µL 20 mM ammonium phosphate in LC-MS-grade water prior to analysis.

***Acquisition parameters***

LC-MS analysis was performed on a Vanquish ultra-high-performance liquid chromatography system coupled to a Q Exactive orbitrap mass spectrometer by a HESI-II electrospray ionization probe (Thermo). External mass calibration was performed weekly. Metabolite samples (2.5 µL) were separated using a ZIC-pHILIC stationary phase (2.1 × 150 mm, 5 µm) (Merck). The autosampler temperature was 4 °C and the column compartment was maintained at 25 °C. Mobile phase A was 20 mM ammonium carbonate, 0.1% ammonium hydroxide, and 5 µM medronic acid. Mobile phase B was acetonitrile. The flow rate was 0.1 mL/min. Solvent was introduced to the mass spectrometer *via* electrospray ionization with the following source parameters: sheath gas 40, auxiliary gas 15, sweep gas 1, spray voltage +3.0 kV for positive mode and -3.1 kV for negative mode, capillary temperature 275 °C, S-lens RF level 40, and probe temperature 350 °C. Data were acquired and peaks integrated using TraceFinder 4.1 (Thermo).

***Metabolomic profiling***

For metabolomic profiling, the following mobile phase gradient was used: 0 min, 80% B; 20 min, 20% B; 20.5 min, 80% B; 28 min, 80% B; 42 min, 80% B. The mass spectrometer was operated in polarity switching full scan mode from 70-1000 m/*z*. Resolution was set to 70,000 and the AGC target was $1\times10^6$ ions. Peak identifications were based on an in-house library of authentic metabolite standards previously analyzed utilizing this method. For metabolomics studies, pooled quality control (QC) samples were injected at the beginning, end, and between every four samples of the run. Raw peak areas for each metabolite were corrected for instrument drift using a cubic spline model of QC peak areas. Low quality features were removed on the basis of a relative standard deviation greater than 0.2 in the QC samples and a dispersion ratio greater than 0.4 *(77)*. Missing values were imputed using random forest. Sample peak areas were normalized using probabilistic quotient normalization *(78)*. Differentially regulated metabolites were identified using limma *(79)*. Exploratory data analysis and review of PCA plots suggested orthogonal effects of TGFβ and MCT inhibitor treatment. For this reason, differentially regulated metabolites were determined based on the main effects of TGFβ or MCT inhibitor. Metabolite set enrichment analysis was performed using the fgsea package *(80)* with KEGG metabolite pathways *(81)*.

***Stable isotope quantification***

Metabolites were measured using the following mobile phase gradient: 0 min, 80% B; 5 min, 80% B; 30 min, 20% B; 31 min, 80% B; 42 min, 80% B. The mass spectrometer was operated in selected ion monitoring mode with an m/*z* window width of 9.0 centered at 1.003355-times half the number of carbon atoms in the target metabolite. The resolution was set at 70,000 and AGC target was $1\times10^5$ ions. Peak areas were corrected for quadrupole bias as in Kim *et al. (*82*)*. Raw mass isotopomer distributions were corrected for natural isotope abundance using a custom R package (mzrtools, github.com/oldhamlab/mzrtools) employing the method of Fernandez, *et al. (83)*.

**RNA-seq transcriptomics**

RNA was collected from LFs treated with TGFβ and MCT inhibitors as described above. Four biological replicates were analyzed. Library construction and sequencing was performed by BGI Genomics using 100 bp paired end analysis and a read depth of 50 M reads per sample. Sequences were mapped to the human GRCh38 primary assembly and counts summarized using salmon *(*84*)*. Differentially expressed genes were identified using DESeq2 *(85)*. Gene set enrichment was performed using the fgsea R package *(80)*.

**Pyridine dinucleotide measurements**

Cellular $NAD^+$ and NADH were measured using an enzymatic fluorometric cycling assay as previously described *(43, 86)*. Samples for $NADP^+$ and NADPH quantification were similarly prepared and analyzed using the luminescence-based NADP/NADPH-Glo assay (Promega).

**Reactive oxygen species**

Intracellular and mitochondrial ROS were measured using the fluorescent probes CellROX Deep Red (1 µM) and MitoSOX Green (2 µM), respectively. Mitochondria were also labeled with MitoTracker (200 nM). Briefly, lung fibroblasts were seeded in a 24-well plate and serum starved for 24 h prior to treatment with TGFβ, with or without MCT inhibitors, for 3 hours. This was followed by staining with the specified probes for 30 min at 37 °C. The cells were then harvested and analyzed using the CytoFLEX flow cytometry analyzer (Beckman Coulter). The mean fluorescence intensity of DAPI-negative cells was analyzed using FlowJo V10.8.1.

**Multi-isotope imaging mass spectrometry**

For multi-isotope imaging mass spectrometry, mice received five intraperitoneal doses of 250 µL of [$^2H_7$]-glucose or [U-$^{13}C_6$]-glucose (200 mg/mL) and [$^{15}N$]-proline (20 mg/mL) every 12 h prior to euthanasia on either day 21 or day 14 after bleomycin. Perfused and inflated lungs were fixed with formaldehyde/glutaraldehyde 2.5% in sodium cacodylate buffer, pH 7.4, embedded in EPON, section to 0.5 µm, mounted on silicon wafers, and gold coated. Sections were analyzed with the NanoSIMS 50L (CAMECA) housed in the Brigham and Women's Hospital Center for NanoImaging, using previously developed analytical protocols *(63, 87)*. The instrument was tuned to simultaneously acquire images of $^{12}C_2{}^1H$, $^{12}C^{14}N$, $^{12}C_2{}^2H$, and $^{12}C^{15}N$ secondary ions. Spatial distribution visualization and quantification of $^{15}N$-proline and $^2H$-glucose labeling was derived from $^{12}C^{15}N/^{12}C^{14}N$ and

$^{12}C_2{}^2H/^{12}C_2{}^1H$ ratios, respectively, using an open source plugin to ImageJ, OpenMIMS 3.0 (github.com/BWHCNI/OpenMIMS). Ratio images are presented visually as hue saturation intensity images, with the blue lower bound of the scale set to the natural background ratio and the red upper bound of the scale set such that within tissue labeling differences are visually apparent. Changes to the scaling modify the visual appearance but not affect the quantitative data. Quantification of isotopic enrichment was based on a pixel ratio analysis of several MIMS images acquired per animal. Specifically, image masks were generated by Otsu thresholding the $^{12}C^{14}N$ to exclude background regions. The ratio of “heavy” to “light” ion intensities in tissue regions was calculated. Pixel ratios equal to zero were excluded. The average of the remaining pixel ratios was then averaged to provide a mean tissue isotope enrichment value for the image.

**VB253**

VB253 was developed by Vettore, LLC as a more potent small molecule inhibitor of MCT4. Experiments involving VB253 were conducted by several contract research organizations under the supervision of Vettore, LLC, as described in detail below. These experiments were completed independently of the other studies presented in the manuscript and offer independent validation of the antifibrotic effects of MCT4 inhibition in cells and mice.

All cell culture experiments were conducted by Charles River Laboratories (Wilmington, MA, USA). Lung fibroblasts from three independent IPF donors were seeded at a density of 750 cells/well in 384-well plates. Fibroblasts were subsequently treated with medium with 0.1% DMSO (vehicle control), 1.25 ng/ml TGFβ alone, 1.25 ng/ml TGFβ in the presence of 1 µM SB525334 (positive control), or 1.25 ng/ml TGFβ with 8-point concentration response curves (CRC) in biologic duplicate with semi-log dilutions of nintedanib or VB253. The top concentration tested was 10 µM. Cells were fixed with 4% paraformaldehyde 72 h post treatment with TGFβ. The principal readout was immunofluorescence staining to for α-SMA and DAPI to quantify the proportion of α-SMA-positive cells.

SMAD3 nuclear translocation was evaluated in lung fibroblasts from one IPF donor. Cells were plated on Purecol-coated 96-well plates. Unstimulated (medium only) and stimulated cells (0.5 ng/ml TGFβ) were treated with 0.1% DMSO (vehicle control), 1 µM SB525334 (positive control), nintedanib in 8-point CRC, ALK5 inhibitor (SB525334) in 8-point CRC, and VB253 in a 16-point CRC, with a maximum concentration of 30 µM. All cells were fixed 2 h after treatment. The readout was immunofluorescence staining for pSmad3 and DAPI.

The *in vivo* therapeutic evaluation of VB253 in bleomycin-induced pulmonary fibrosis in both young and aged mice was conducted by Aragen Biosciences (Morgan Hill, CA, USA). In two separate experiments, C57Bl/6 mice, aged either 6-7 weeks or 68-70 weeks, were obtained from Jackson Labs (Bar Harbor, ME, USA). Under anesthesia, mice were administered either PBS vehicle (N = 10 per experiment) or 1.5 U/kg bleomycin in PBS intratracheally. After 7 days, mice treated with bleomycin were randomized to receive either vehicle (0.5% methylcellulose in saline) (N = 10), VB253 dissolved in 0.5% methylcellulose (3 mg/kg BID), Pirfenidone (Glentham Life Sciences Lid. Cat #GP4948, 100 mg/kg BID), or Nintedinib (Axon MedChem Cat#2648, 50 mg/kg, daily) (N = 10 per group). Mice injected with saline, rather than bleomycin, were administered vehicle. Treatments were administered daily until the experiment was terminated 21 days after bleomycin instillation, approximately 2-4 h after the final treatment dose. Whole-body plethysmography was performed on day 20 using Buxco WBP instrument system measuring Enhanced Pause (Penh) and respiratory rate on unrestrained animals. Upon experiment termination, lungs were inflated with 10% neutral buffered formalin (NBF) and fixed in 10% NBF for histology.

Histology, imaging, and pathologic analyses were performed by HistoTox Labs (Boulder, CO, USA). Multiple slides per block were sectioned at 5 µm and stained with hematoxylin and eosin (H&E) or by routine immunohistochemistry (IHC) methods for α-SMA. H&E-stained glass slides were evaluated using light microscopy by an ACVP board-certified veterinary pathologist. Lung sections were scored according to the modified Ashcroft scale. Briefly, scores for five representative 200× microscopic fields per sample were averaged to obtain a mean score for each animal. α-SMA-stained glass slides were scanned using an Aperio AT2 whole slide scanner. Whole slide images were annotated to delineate regions of interest (ROI). Exclusions were applied to remove tissue artifacts (folds, tears, non-specific staining), large airways, and blood vessels. Visiopharm (VIS) image analysis software, using imaging filters to separate positive staining from counterstaining and background were applied to quantify α-SMA signal within each sample. Differences in expression levels between samples and treatments were evaluated by calculating the percent positive detection of cells or area within the viable lung tissue.

### FIGURES

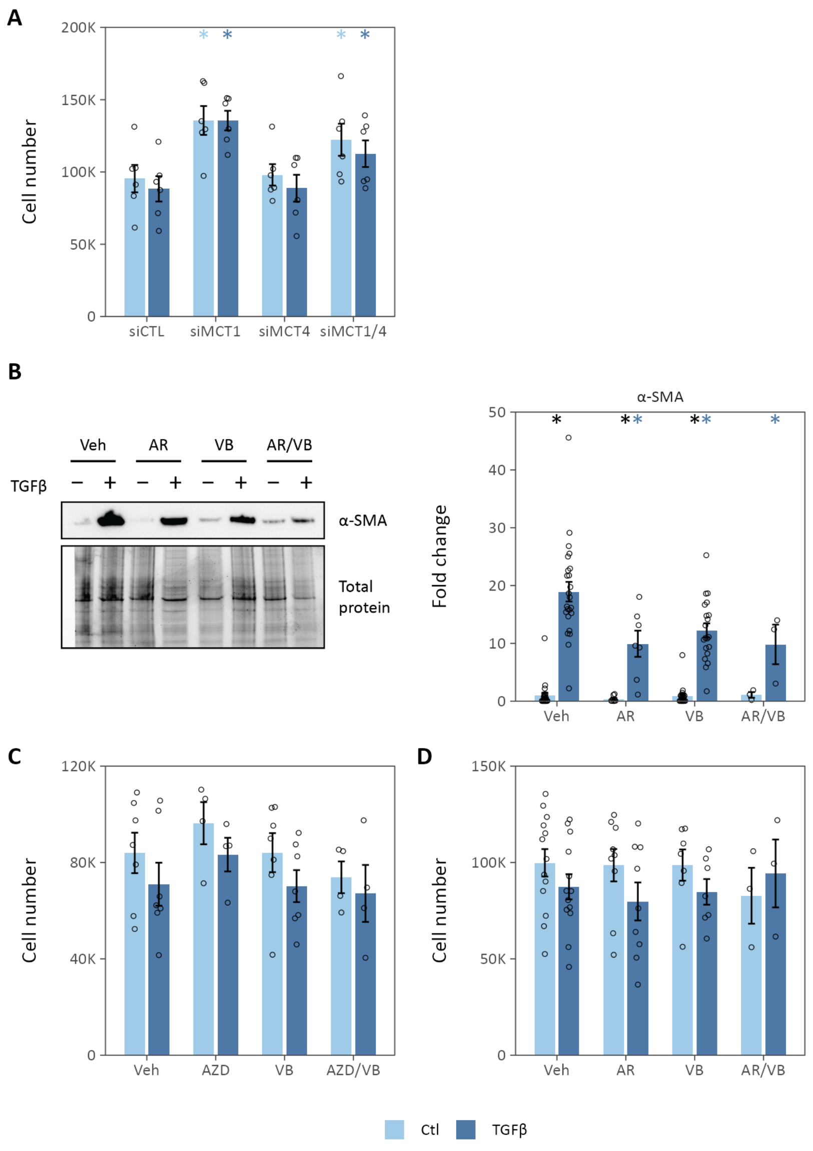


**Fig. S1. Lactate transport inhibition decreases myofibroblast differentiation *in vitro*.** (**A**) Cell count following siRNA-mediated MCT knockdown. (**B**) Small molecule inhibitors of MCT1/2 (AR-C155858, AR) or MCT4 (VB124, VB) decrease TGFβ-stimulated α-SMA expression in normal human lung fibroblasts. (**C**-**D**) Cell counts following pharmacologic inhibition of lactate transporters. Summary data are mean ± SEM (* p-value-value < 0.05; *black* compares TGFβ to control within a given treatment, *colored* compares the treatment effect to control for a given condition).

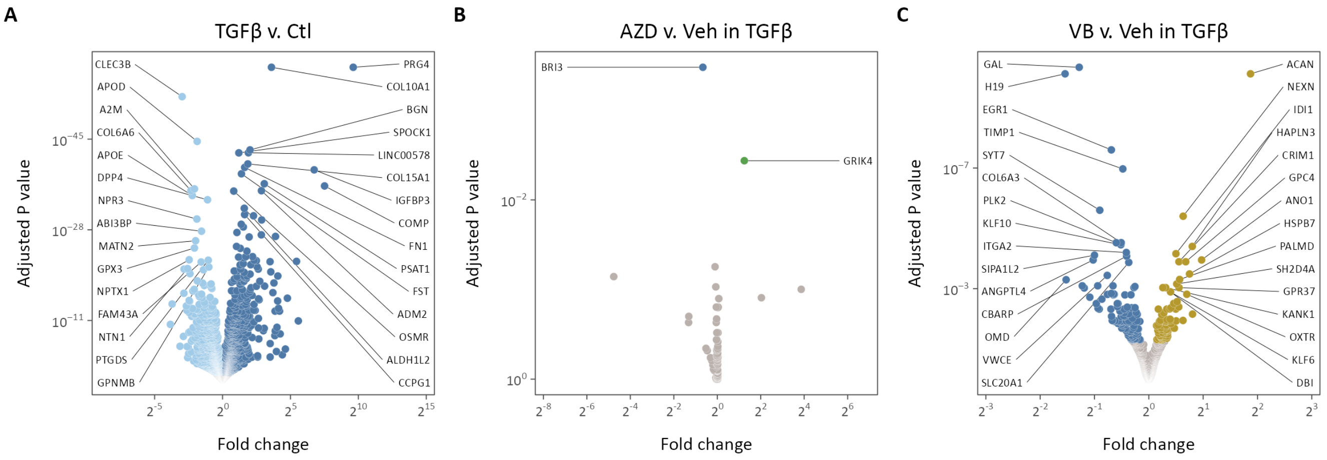


**Fig. S2. Lactate transporter inhibition decreases pro-fibrotic gene transcription.** (**A**-**C**) Volcano plots of differentially expressed genes in TGFβ-treated cells compared to control (A), AZD3965-treated cells compared to vehicle (Veh) (B), and VB124-treated cells compared to Veh. Significantly differentially expressed genes are highlighted (adjusted p-value < 0.05), the top 15 up- and down-regulated of which are labeled.

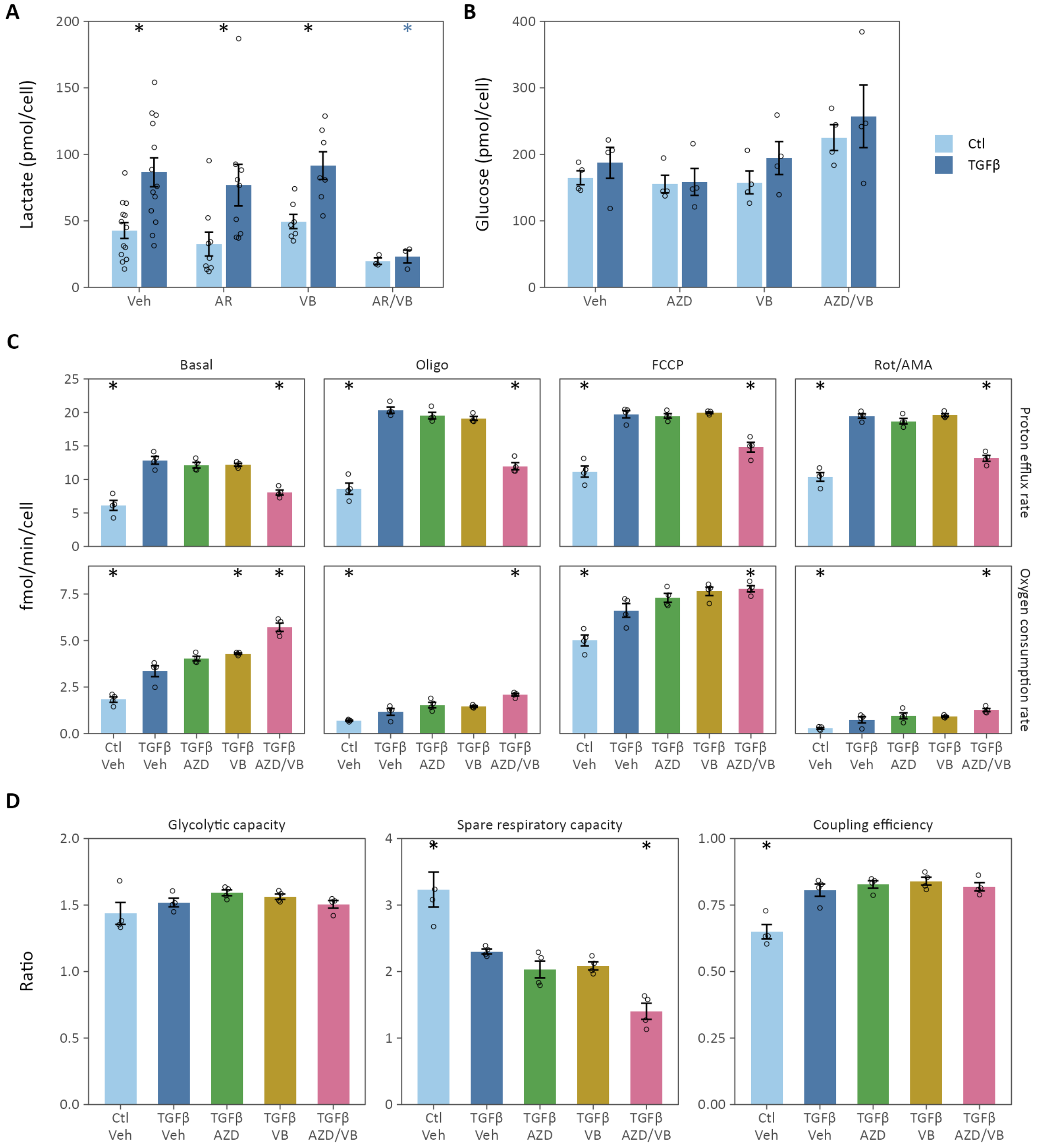


**Fig. S3. Lactate transport inhibition alters cellular bioenergetics.** (**A**) Extracellular lactate was determined 48 h following TGFβ in the presence of MCT1/2 inhibitor AR-C155858 (AR), MCT4 inhibitor VB124 (VB), or both (N = 3-13 biological replicates, * p-value < 0.05, *black* compares TGFβ *v.* Ctl, *colored* compares Drug *v.* Veh). (**B**) Extracellular glucose was determined 48 h following TGFβ in the presence of MCT1 inhibitor AZD3965 (AZD), VB, or both (N = 5 biological replicates). (**C**) Summary data from Seahorse extracellular flux analysis. (**D**) Cell bioenergetic stress testing results from Seahorse analysis. Glycolytic capacity measures the increase in proton efflux rate following oligomycin treatment. Spare respiratory capacity measures the increase in oxygen consumption rate (OCR) following FCCP treatment compared to basal OCR. Coupling efficiency measures the proportion of basal OCR that contributes to ATP production (*i.e.,* the percentage inhibited by oligomycin). The spare respiratory capacity decreases following MCT inhibition, indicating a larger fraction of total respiratory capacity was utilized in the basal state in cells treated with MCT inhibitors (N = 4 biological replicates, * p-value < 0.05 compared to TGFβ/Veh). Summary data are mean ± SEM.

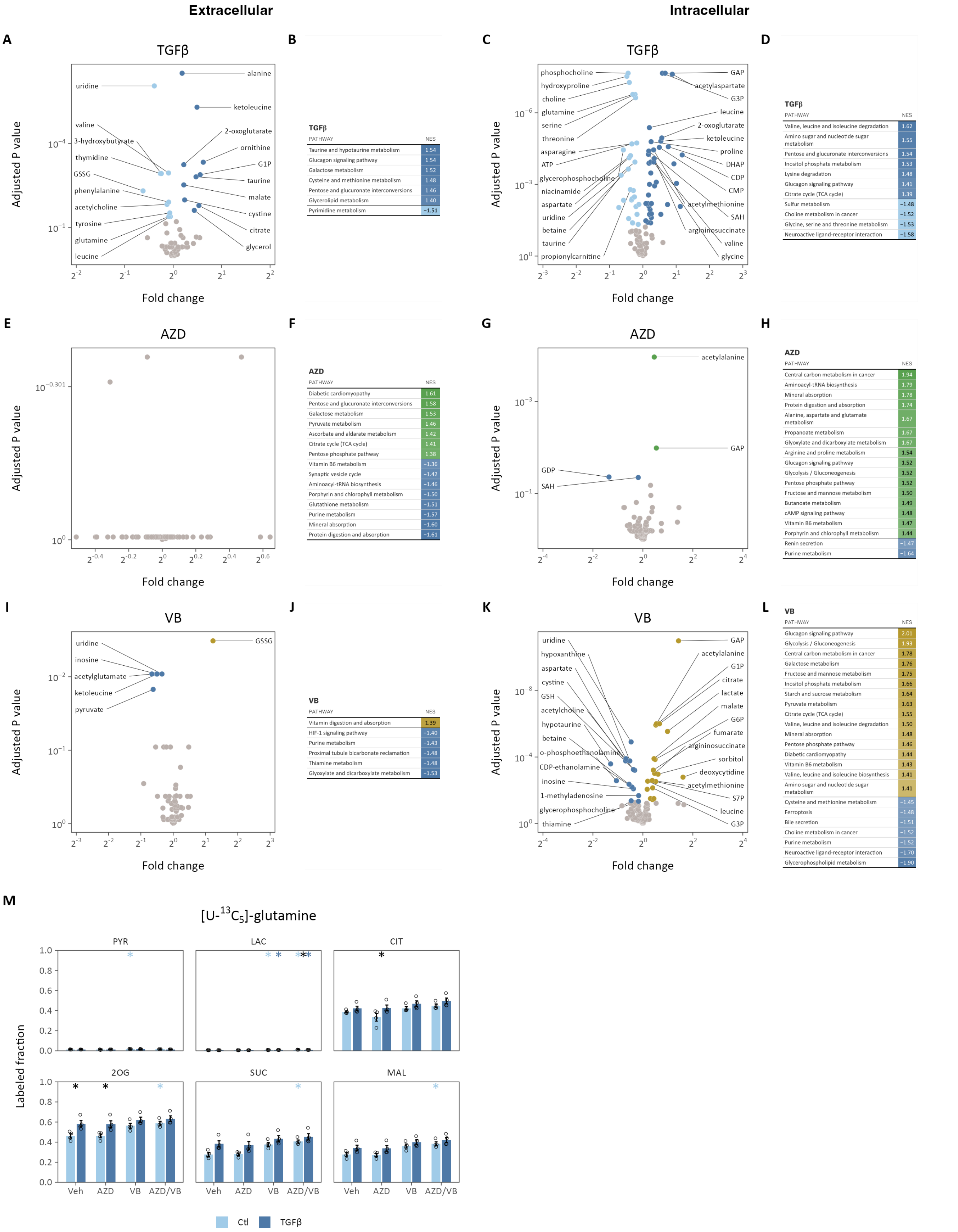


**B**

| TGFβ PATHWAY | NES |
|---|---|
| Taurine and hypotaurine metabolism | 1.54 |
| Glucagon signaling pathway | 1.54 |
| Galactose metabolism | 1.52 |
| Cysteine and methionine metabolism | 1.48 |
| Pentose and glucuronate interconversions | 1.46 |
| Glycerolipid metabolism | 1.40 |
| Pyrimidine metabolism | −1.51 |

**D**

| TGFβ PATHWAY | NES |
|---|---|
| Valine, leucine and isoleucine degradation | 1.62 |
| Amino sugar and nucleotide sugar metabolism | 1.55 |
| Pentose and glucuronate interconversions | 1.54 |
| Inositol phosphate metabolism | 1.53 |
| Lysine degradation | 1.48 |
| Glucagon signaling pathway | 1.41 |
| Citrate cycle (TCA cycle) | 1.39 |
| Sulfur metabolism | −1.48 |
| Choline metabolism in cancer | −1.52 |
| Glycine, serine and threonine metabolism | −1.53 |
| Neuroactive ligand-receptor interaction | −1.58 |

**F**

| AZD PATHWAY | NES |
|---|---|
| Diabetic cardiomyopathy | 1.61 |
| Pentose and glucuronate interconversions | 1.58 |
| Galactose metabolism | 1.53 |
| Pyruvate metabolism | 1.46 |
| Ascorbate and aldarate metabolism | 1.42 |
| Citrate cycle (TCA cycle) | 1.41 |
| Pentose phosphate pathway | 1.38 |
| Vitamin B6 metabolism | −1.36 |
| Synaptic vesicle cycle | −1.42 |
| Aminoacyl-tRNA biosynthesis | −1.46 |
| Porphyrin and chlorophyll metabolism | −1.50 |
| Glutathione metabolism | −1.51 |
| Purine metabolism | −1.57 |
| Mineral absorption | −1.60 |
| Protein digestion and absorption | −1.61 |

**H**

| AZD PATHWAY | NES |
|---|---|
| Central carbon metabolism in cancer | 1.94 |
| Aminoacyl-tRNA biosynthesis | 1.79 |
| Mineral absorption | 1.78 |
| Protein digestion and absorption | 1.74 |
| Alanine, aspartate and glutamate metabolism | 1.67 |
| Propanoate metabolism | 1.67 |
| Glyoxylate and dicarboxylate metabolism | 1.67 |
| Arginine and proline metabolism | 1.54 |
| Glucagon signaling pathway | 1.52 |
| Glycolysis / Gluconeogenesis | 1.52 |
| Pentose phosphate pathway | 1.52 |
| Fructose and mannose metabolism | 1.50 |
| Butanoate metabolism | 1.49 |
| cAMP signaling pathway | 1.48 |
| Vitamin B6 metabolism | 1.47 |
| Porphyrin and chlorophyll metabolism | 1.44 |
| Renin secretion | −1.47 |
| Purine metabolism | −1.64 |

**J**

| VB PATHWAY | NES |
|---|---|
| Vitamin digestion and absorption | 1.39 |
| HIF-1 signaling pathway | −1.40 |
| Purine metabolism | −1.43 |
| Proximal tubule bicarbonate reclamation | −1.48 |
| Thiamine metabolism | −1.48 |
| Glyoxylate and dicarboxylate metabolism | −1.53 |

**L**

| VB PATHWAY | NES |
|---|---|
| Glucagon signaling pathway | 2.01 |
| Glycolysis / Gluconeogenesis | 1.93 |
| Central carbon metabolism in cancer | 1.78 |
| Galactose metabolism | 1.76 |
| Fructose and mannose metabolism | 1.75 |
| Inositol phosphate metabolism | 1.66 |
| Starch and sucrose metabolism | 1.64 |
| Pyruvate metabolism | 1.63 |
| Citrate cycle (TCA cycle) | 1.55 |
| Valine, leucine and isoleucine degradation | 1.50 |
| Mineral absorption | 1.48 |
| Pentose phosphate pathway | 1.46 |
| Diabetic cardiomyopathy | 1.44 |
| Vitamin B6 metabolism | 1.43 |
| Valine, leucine and isoleucine biosynthesis | 1.41 |
| Amino sugar and nucleotide sugar metabolism | 1.41 |
| Cysteine and methionine metabolism | −1.45 |
| Ferroptosis | −1.48 |
| Bile secretion | −1.51 |
| Choline metabolism in cancer | −1.52 |
| Purine metabolism | −1.52 |
| Neuroactive ligand-receptor interaction | −1.70 |
| Glycerophospholipid metabolism | −1.90 |

**Fig. S4. Metabolomic profiling of lung fibroblasts treated with lactate transport inhibitors.** (**A**-**D**) Extracellular (A-B) and intracellular (C-D) metabolomic profile and metabolite set enrichment analyses of lung fibroblasts treated with TGFβ. (**E-H**) Extracellular (E-F) and intracellular (G-H) metabolomic profile and metabolite set enrichment analysis of AZD3965 treatment. (**I-L**) Extracellular (I-J) and intracellular (K-L) metabolomic profile and metabolite set enrichment analysis of VB124 treatment. Differentially regulated metabolites are colored (adjusted p-value < 0.1), the top 10 up- and down-regulated of which are labeled. KEGG pathways significantly enriched (adjusted p-value < 0.1) with metabolites differentially regulated by TGFβ treatment or MCT inhibition ordered by normalized enrichment score (NES). Positive NES indicates enrichment in treated cells while negative NES indicates enrichment in Vehicle-treated cells. (**M**) Labeled fraction of intracellular metabolites pyruvate (PYR), lactate (LAC), citrate (CIT), 2-oxoglutarate (2OG), succinate (SUC), and malate (MAL) following treatment with [U-$^{13}C_5$]-glutamine (N = 4 biological replicates, * adjusted p-value < 0.05, *black* compares TGFβ *v.* Ctl for a given treatment, *colored* compares treatment *v.* vehicle for the indicated condition). Summary data are mean ± SEM.

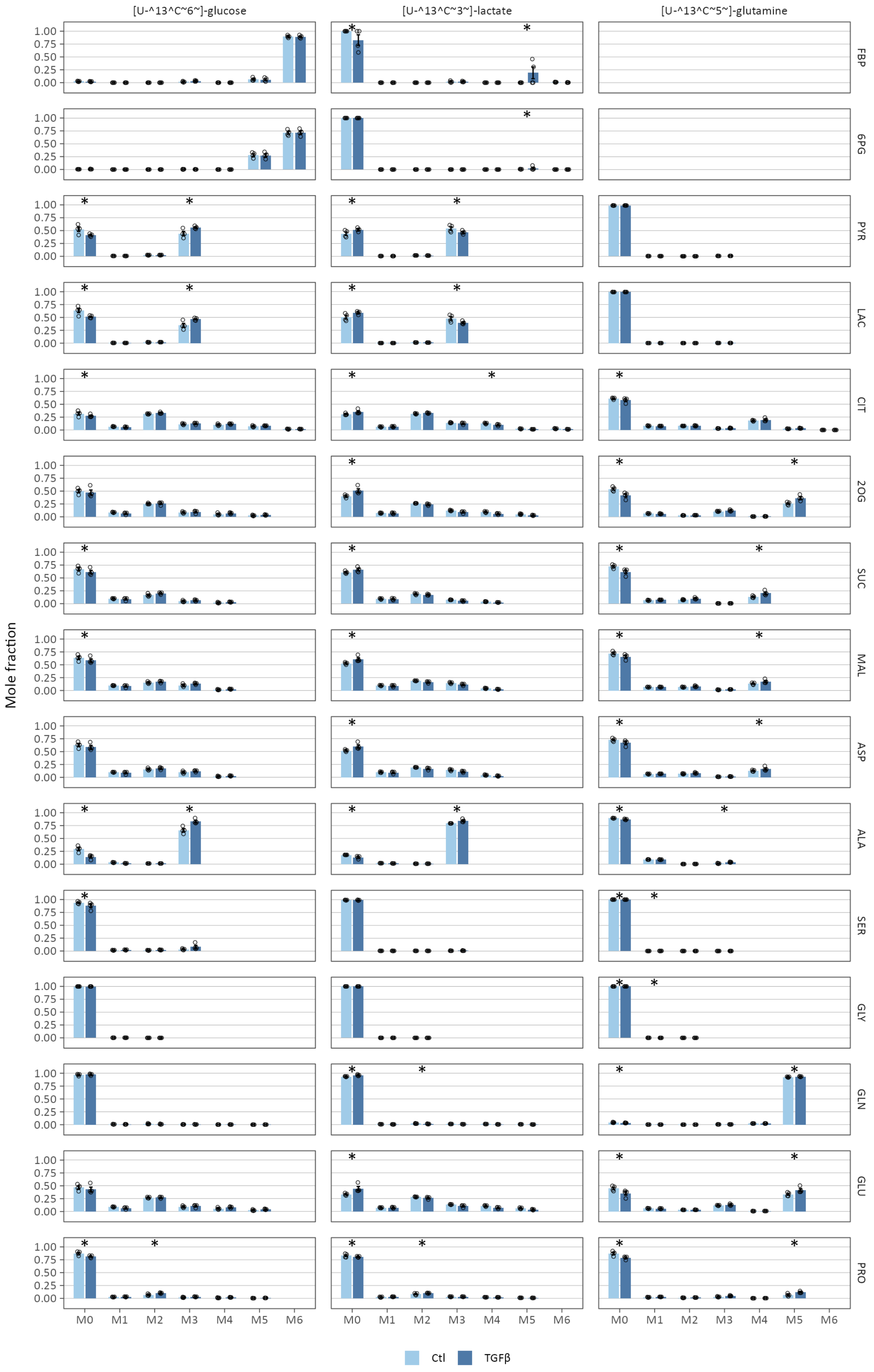
[U-^13^C~6~]-glucose
[U-^13^C~3~]-lactate
[U-^13^C~5~]-glutamine
FBP
6PG
PYR
LAC
CIT
2OG
SUC
MAL
ASP
ALA
SER
GLY
GLN
GLU
PRO
Mole fraction
1.00
0.75
0.50
0.25
0.00
M0
M1
M2
M3
M4
M5
M6
Ctl
TGFβ

**Fig. S5. Stable isotope tracing in lung fibroblasts treated with TGFβ.** Lung fibroblasts were cultured with stable isotopes of glucose, lactate, and glutamine during treatment with TGFβ (*columns*). Mass isotope distributions were determined by LC-MS for key metabolites in central carbon metabolic pathways (*rows*). Significant differences in isotopic enrichment are indicated (* adjusted p-value < 0.05). Summary data are mean ± SEM (N = 4 biological replicates).

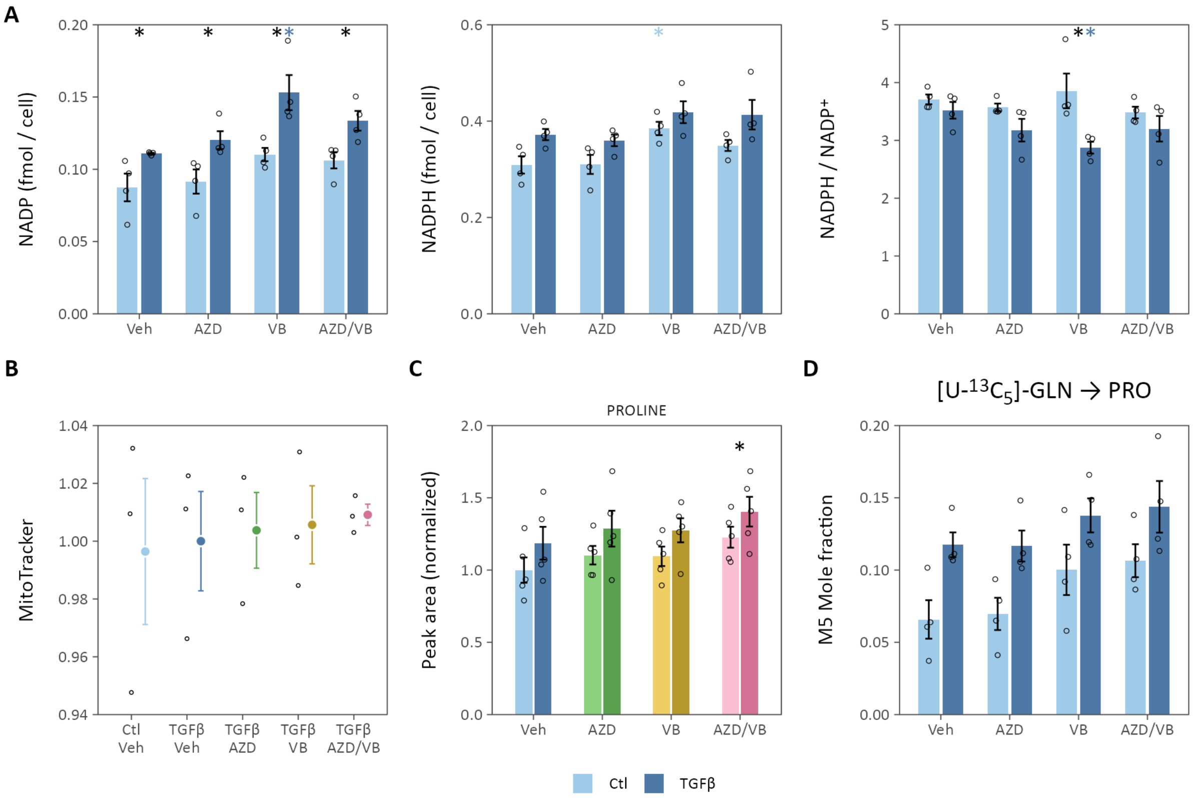


**Fig. S6. Lactate transport inhibition activates antioxidant defenses.** (**A**) The cellular redox couple NADPH/NADP⁺ was determined by enzymatic cycling assay (N = 4 biological replicates; * adjusted p-value < 0.05; *black* compares TGFβ to control within a given treatment, *colored* compares the treatment effect to control for a given condition). (**B**) MitoTracker fluorescence was measured as a marker of mitochondrial mass (N = 3 biological replicates; * adjusted p-value < 0.05 compared to TGFβ-treated cells). (**C**) Proline peak areas determined by LC-MS (N = 5 biological replicates, * adjusted p-value < 0.05 for the main treatment effect). (**D**) Fraction of the M5 proline isotope following labeling with [U-$^{13}C_5$]-glutamine (N = 4 biological replicates). Summary data are mean ± SEM.

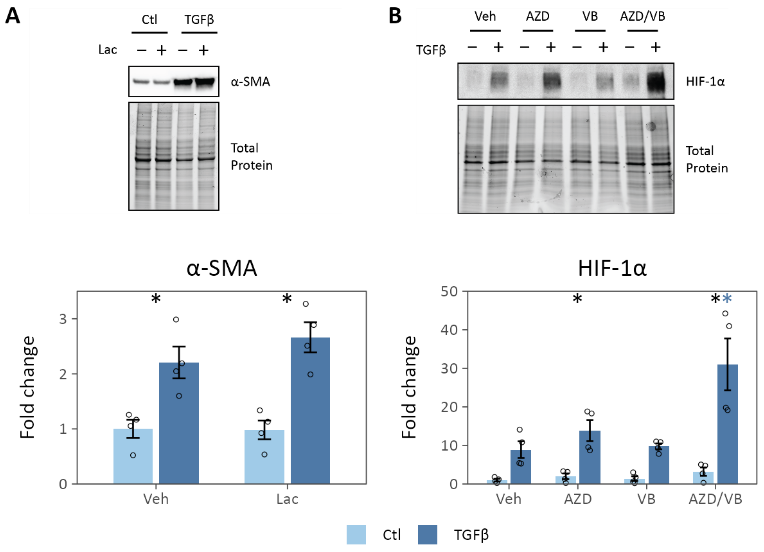


**Fig. S7. Mechanisms of lactate signaling.** (**A**) Exogenous lactate (10 mM) does not ameliorate TGFβ-stimulated α-SMA expression. (**B**) Lactate transport inhibition does not attenuate HIF-1α stabilization 6 h after TGFβ treatment. Summary data are mean ± SEM. (N = 4 biological replicates, * adjusted p-value < 0.05, *black* compares TGFβ *v.* Ctl for a given treatment, *colored* compares treatment *v.* vehicle for the indicated condition).

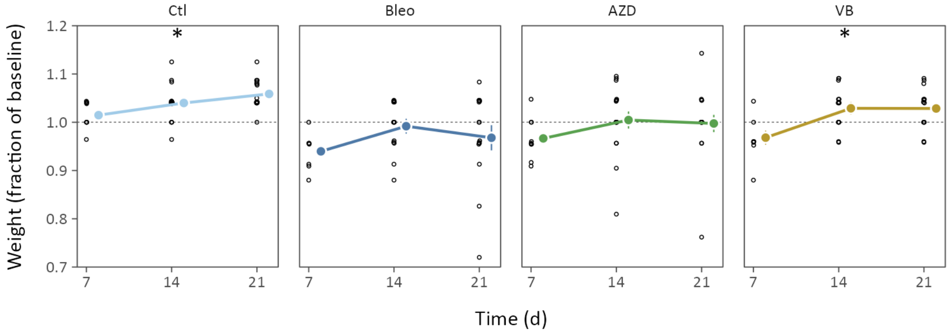


**Fig. S8. Lactate transport inhibition improves weight gain following bleomycin.** Following bleomycin, mice were weighed weekly. Data points show individual mice, summary statistics show the mean ± SEM, * adjusted p-value-value < 0.05 for the overall treatment effect compared to bleomycin-treated (Bleo) vehicle control.

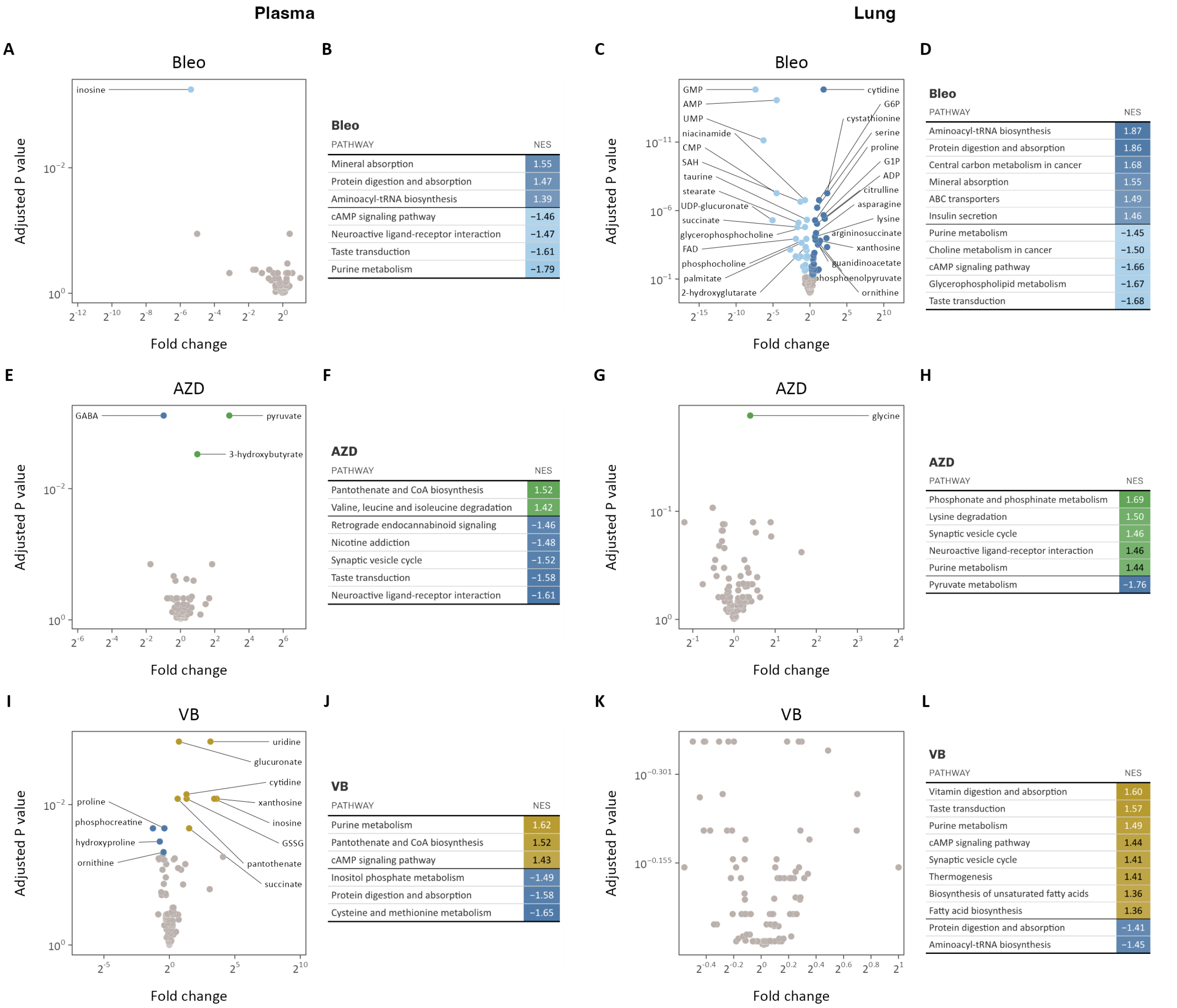


**Bleo** (B)

| PATHWAY | NES |
|---|---|
| Mineral absorption | 1.55 |
| Protein digestion and absorption | 1.47 |
| Aminoacyl-tRNA biosynthesis | 1.39 |
| cAMP signaling pathway | −1.46 |
| Neuroactive ligand-receptor interaction | −1.47 |
| Taste transduction | −1.61 |
| Purine metabolism | −1.79 |

**Bleo** (D)

| PATHWAY | NES |
|---|---|
| Aminoacyl-tRNA biosynthesis | 1.87 |
| Protein digestion and absorption | 1.86 |
| Central carbon metabolism in cancer | 1.68 |
| Mineral absorption | 1.55 |
| ABC transporters | 1.49 |
| Insulin secretion | 1.46 |
| Purine metabolism | −1.45 |
| Choline metabolism in cancer | −1.50 |
| cAMP signaling pathway | −1.66 |
| Glycerophospholipid metabolism | −1.67 |
| Taste transduction | −1.68 |

**AZD** (F)

| PATHWAY | NES |
|---|---|
| Pantothenate and CoA biosynthesis | 1.52 |
| Valine, leucine and isoleucine degradation | 1.42 |
| Retrograde endocannabinoid signaling | −1.46 |
| Nicotine addiction | −1.48 |
| Synaptic vesicle cycle | −1.52 |
| Taste transduction | −1.58 |
| Neuroactive ligand-receptor interaction | −1.61 |

**AZD** (H)

| PATHWAY | NES |
|---|---|
| Phosphonate and phosphinate metabolism | 1.69 |
| Lysine degradation | 1.50 |
| Synaptic vesicle cycle | 1.46 |
| Neuroactive ligand-receptor interaction | 1.46 |
| Purine metabolism | 1.44 |
| Pyruvate metabolism | −1.76 |

**VB** (J)

| PATHWAY | NES |
|---|---|
| Purine metabolism | 1.62 |
| Pantothenate and CoA biosynthesis | 1.52 |
| cAMP signaling pathway | 1.43 |
| Inositol phosphate metabolism | −1.49 |
| Protein digestion and absorption | −1.58 |
| Cysteine and methionine metabolism | −1.65 |

**VB** (L)

| PATHWAY | NES |
|---|---|
| Vitamin digestion and absorption | 1.60 |
| Taste transduction | 1.57 |
| Purine metabolism | 1.49 |
| cAMP signaling pathway | 1.44 |
| Synaptic vesicle cycle | 1.41 |
| Thermogenesis | 1.41 |
| Biosynthesis of unsaturated fatty acids | 1.36 |
| Fatty acid biosynthesis | 1.36 |
| Protein digestion and absorption | −1.41 |
| Aminoacyl-tRNA biosynthesis | −1.45 |

**Fig. S9. Metabolic reprogramming by lactate transport inhibition *in vivo*.** (**A**-**D**) Plasma (A-B) and lung (C-D) metabolomic profile and metabolite set enrichment analyses in bleomycin-treated mice. (**E**-**H**) Plasma (E-F) and lung (G-H) metabolomic profile and metabolite set enrichment analysis of AZD3965 treatment in bleomycin-treated mice. (**I**-**L**) Plasma (I-J) and lung (K-L) metabolomic profile and metabolite set enrichment analysis of VB124 treatment in bleomycin-treated mice. Differentially regulated metabolites are colored (adjusted p-value < 0.1), the top 10 up- and down-regulated of which are labeled. KEGG pathways significantly enriched (adjusted p-value < 0.1) with metabolites differentially regulated by bleomycin treatment or MCT inhibition ordered by normalized enrichment score (NES). Positive NES indicate enrichment in treated mice while negative NES indicate relative enrichment in control or untreated mice.

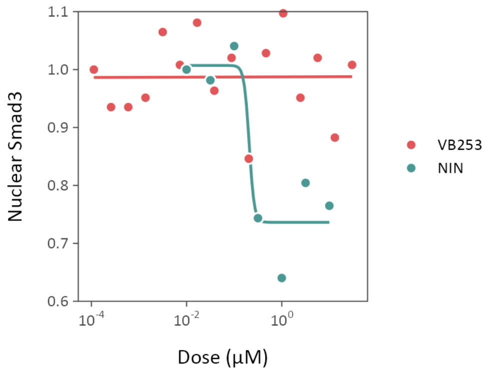


**Fig. S10. VB253 does not attenuate canonical TGFβ signaling.** Upon phosphorylation by the TGFβ receptor, Smad3 relocates to the nucleus. Dose-response profile of VB253 compared to nintedanib (NIN), demonstrates that VB253 does not inhibit canonical Smad3 signaling pathways.
